# Individual brain activity patterns during task are predicted by distinct resting-state networks that may reflect local neurobiological features

**DOI:** 10.1101/2024.11.13.621472

**Authors:** Robert Scholz, R. Austin Benn, Victoria Shevchenko, Ulysse Klatzmann, Wei Wei, Francesco Alberti, Rocco Chiou, Xi-Han Zhang, Robert Leech, Jonathan Smallwood, Daniel S. Margulies

**Affiliations:** Cognitive Neuroanatomy Lab, Université Paris Cité, INCC UMR 8002, CNRS, Paris, France; Wellcome Centre for Integrative Neuroimaging, FMRIB, Nuffield Department of Clinical Neurosciences, University of Oxford, UK; Wilhelm Wundt Institute for Psychology, Leipzig University, Leipzig, Germany; Max Planck School of Cognition, Leipzig, Germany; Université Paris-Saclay, Inria, CEA, Palaiseau, France; School of Psychology, University of Surrey, Surrey, UK; Department of Psychology, Yale University, New Haven, CT, USA; Department of Neuroimaging, King’s College London, UK; Department of Psychology, Queen’s University, Kingston, Canada

## Abstract

Understanding how individual cortical features shape functional brain organization offers a promising framework for examining the principles of cognitive specialization in the human brain. This study explores the relationship between various cortical characteristics—i.e resting-state functional connectivity, structural connectivity, microstructure, morphology, and geometry—and the layout of task-specific functional activations. We employ linear models to predict the functional layout of the cortex at the individual level from each of these feature modalities. Our findings demonstrate that resting-state component loadings predict individual task activations, consistently across hemispheres and independent datasets. Whereas the first few components provide a common space for functional activations across tasks, predictive higher-order component loadings demonstrated task-specificity. Cortical microstructure/morphology was notably predictive of activation strength in the occipital cortex, highlighting its relevance for cortical functional specialization. By relating resting state components to a set of reference maps of cortical organization, we identify associations that suggest possible neurobiological underpinnings of specific cognitive functions. The remaining feature modalities were only predictive of group-level functional activations. These results advance our understanding of how distinct cortical features may contribute to functional specialization, guiding future inquiry into the organization of cognitive functions on the cortex.

## Introduction

The layout of functions in the cerebral cortex is reflected in unique features of its constituent areas. Specifically, cortical microstructure, reflecting local features such as gene expression, cell types and laminar differentiation may give rise to specialized computational properties. A complementary view is that connectivity between cortical areas drives the unique functional roles that each area subsumes. Further accounts have highlighted the role that cortical geometry — possibly as a proxy for local connectivity — may play in the global layout of functions (Griffins 1995; Plaut 2002; Goldberg 1995; Saadon-Grosmann 2015; Margulies, 2016; Wang, 2023; Pang 2023).

These ideas can be rendered more concrete based on the example of language. The brain’s language network is centered around two core regions - historically referred to as Broca’s and Wernicke’s area. These areas can be characterized by their specific cytoarchitectural composition and gene expression (Amunts, 2007, Unger, 2021), connectivity (including their interconnection through the arcuate fasciculus, see Rilling, 2008) and positioning on the cortex (being relatively distal to sensory areas, possibly at the intersection of processing streams). These characteristics may all be of functional relevance (e.g. Enard, 2002; Reimers-Kipping, 2011; Rilling, 2008). Similarly, other functions may be characterized and constrained by distinct sets of features.

Group-level analyses have been instrumental in revealing relationships between the brain’s macroscale functional organization and specific cortical features. The understanding of brain function can be further enriched by examining individual-level differences. For example, if variations in the spatial distribution of a feature at the individual level are reflected in functional activations, this would suggest a strong influence of that feature in shaping the cortical layout of functional specialization, providing additional evidence for specific theoretical accounts. Previous studies have demonstrated that the individual functional layout can be accurately predicted based on resting-state fMRI defined measures (Langs, 2015; Tavor, 2016; Jones, 2017; Tobyne, 2019; Ngo, 2022; Zheng, 2022; Tik, 2023). In the current study, we use similar methods to probe a wider set of features, and identify those that are strongly predictive of functional specialization in the cortex.

As predictors in our models, we used seven types of cortical features — referred to as *feature modalities* in the remainder of the manuscript — covering connectivity, microstructure/morphology, and geometry. This includes individualized resting-state functional connectivity measures (PCA and ICA component loadings), a measure of structural connectivity (structural connectivity blueprints, Warrington, 2020), and measures of microstructure/morphology (i.e. T1/T2, cortical thickness, curvature and sulcal profile). Additionally, we incorporated three measures of cortical geometry— i.e. distances on the cortical surface to reference landmarks, PCA components of the resulting distance matrix and geometric eigenmodes — all derived from individual cortical surfaces.

Based on these feature modalities, we predict individual functional activations, i.e. task contrast maps, in seven tasks. We then proceed to quantify and interpret the contributions of features within each modality. This setup allows us to assess how a given modality may relate to the emergence of functions at precise cortical locations. By systematically analyzing these relationships, we can discern patterns and potential principles underlying cortical functional organization. We employ these analyses to further our understanding of how biology has managed to ensure that specific functions can be reliably acquired by distinct regions of the human brain based on a limited set of organizational features.

## Methods

Our approach involved predicting individual task contrasts by fitting linear models on various modalities. To evaluate model performance, we compared model predictions to the ground truth. To understand why certain task contrasts tend to be more readily predicted than others, we contextualized the resulting metrics by computing comparable measures (prediction baselines) on a separate test-retest dataset. This quantified the stability of individual task contrast maps across two separate scanning runs, and provides an approximate ceiling for the prediction accuracy that differed from contrast to contrast. We analyzed model parameters to identify the features that drive these predictions, and compared the spatial distribution of the most predictive features to reference brain maps (Figure 1f) to render them more interpretable. We conducted these analyses separately for each hemisphere, and validated our findings in an independent dataset.

**Figure 1.**
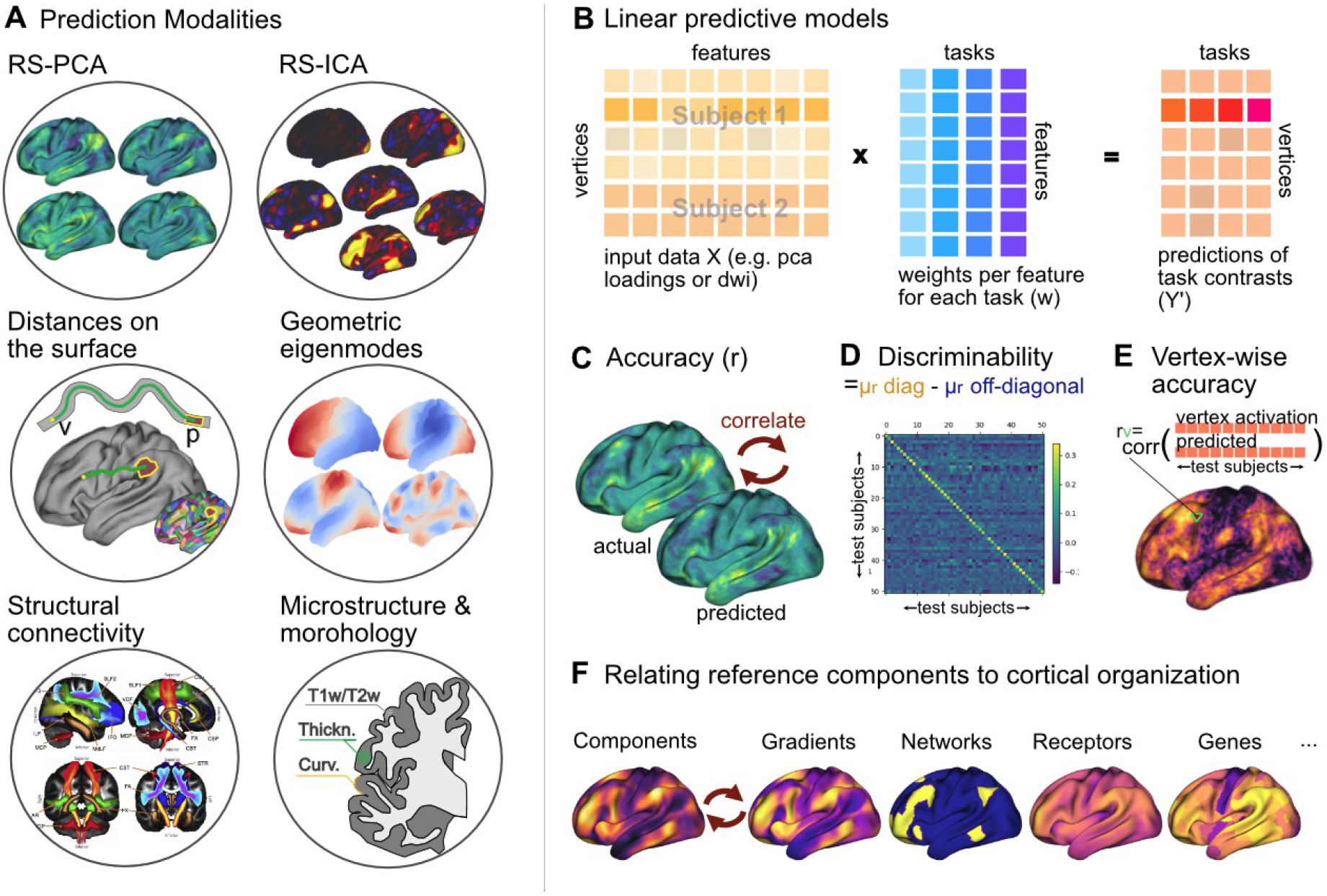
Overview of the prediction model and scores used for the assessment of model performance. **A** Functional and structural measures used as predictors in our models, including resting-state functional connectivity PCA (RS-PCA) and ICA (RSICA) component loadings, distances on the cortical surface (in two varieties, i.e. distances to landmarks, and distance matrix PCA component loadings), geometric eigenmodes of the cortical surface, structural connectivity blueprints (image based on a section of Figure 3 from Warrington, 2020), and cortical microstructure/morphology. **B** Predictors consist of a set of features per vertex. We then fit a linear model taking these features as input to predict all target task contrasts. The same mapping (=weights) is applied at each position (=vertex) on the cortex and across subjects. Given that we constrain this mapping to be agnostic to the precise location on the cortex and specific subject, we can pool vertex-wise feature and task maps from all subjects during model estimation (as indicated in the figure by highlighting specific portions of the features across pertaining to two different subjects). **C-E** Model performance is quantified using three distinct scores. **C** Overall accuracy is estimated by correlating real and predicted task maps. **D** To estimate if prediction results actually capture interindividual differences, we compute a discriminability score. It is calculated as the difference between the average correlation of predicted and actual subjects’ target task maps (values on the diagonal), minus the average correlation of predicted and any other subjects’ target map (off-diagonal). **E** Vertex-wise accuracy is another measure aimed at assessing model performance in predicting individual differences. At each vertex, the series of predicted values for all the subjects is correlated with the real series of values. This gives an approximation of the regional performance of the model. **F** We correlate feature maps that do not have a straightforward biological interpretation (such as resting-state connectivity components) with reference maps (reflecting e.g. resting-state connectivity gradients, functional networks, receptors and gene expression) to identify the reference maps that most closely mirror each feature map. This facilitates the interpretation of what these feature maps may be reflective of and how they could relate to specific task contrast maps.

### Datasets

We used the minimally preprocessed and MSMAll-registered 3T functional magnetic resonance imaging (fMRI) data from the Human Connectome Project (HCP) young adult 1200 subjects data release (Glasser, 2013; Van Essen, 2013). The main dataset consisted of 254 healthy, unrelated HCP subjects (aged 22–35 years, 132 females and 122 males). This subset is based on the 255 HCP subjects used by Pang et al. (Pang, 2023), which represents the largest HCP sample excluding twins or siblings with complete task-evoked and resting-state data. One subject was excluded due to missing preprocessed files. We further used the test-retest subset of HCP - 45 subjects for which functional data acquisition was repeated - to derive model score baselines (see section “prediction baselines”).

### Task maps (prediction targets)

As prediction targets, we used the t-statistic contrast maps from the left hemisphere in fsLR standard surface space at a sampling density of 32k (fsLR32k), excluding vertices that form part of the medial wall. This yielded one map per task contrast and subject, describing the functional activations at all of the 29k left hemispheric vertices. These maps were z-scored across vertices, before being used as prediction targets. We predicted all available contrasts for the seven tasks of the HCP dataset (Barch, 2013), excluding sign-flipped variants of the same contrast, e.g. “Story-Math” vs “Math-Story”. This yielded 47 contrasts in total. For the result visualizations in the main text, we selected the task contrasts with the most reliable interindividual differences in the test-retest dataset for each of the 7 tasks, as assessed by their test-retest discriminability (see baselines subsection and Supplementary Table 1). The full results are available in the supplementary materials.

In general, tasks took 2-8 mins each for completion, and within a task, trials of the same condition were usually presented in blocks. The ***working memory (WM)*** task tested both for category specific representations and working memory. In this task, participants saw stimuli of 4 categories: places, tools, faces and body parts, presented in separate blocks. Depending on the task condition, participants had to respond whenever the current stimulus matched a target given at the beginning of the block (0-back) or whenever it matched the stimulus two presentations before (2-back). During the ***gambling task***, participants had to guess whether the number of a hidden card was above or below 5, winning either 1$ if correct (reward), losing $0.5 if wrong (punish), or nothing if the card was exactly 5 (neutral). During the ***motor task***, participants were visually cued to perform specific movements using different body parts (tapping left or right finger, squeezing left or right toe, or moving their tongue) during a 12 second post-cue period. In the ***language task***, participants either listened to short auditory stories or math problems, each followed by a question (regarding the stories’ topic, or the math problems’ solution) and a period in which the subjects had to choose between two possible answers through button press. In the ***social cognition task***, participants watched short videos in which a set of moving objects (squares, circles, and triangles) either seemed to interact socially (“mentally”), or in which the object movements seemed to be random. Participants reported the interaction type (social, random, unsure) by pressing a button. In the **emotion task**, participants had to indicate whether a presented face or shape matches a concurrently presented target, with the faces having a fearful or angry expression. Lastly, in the **relational processing task** trials, participants had to decide whether two pairs of objects differed along the same dimension (either shape or texture) or not (“relational processing” condition). Alternatively, they had to decide whether a single object matched either of two separate objects in a specific dimension that was cued by a word (e.g. “shape”) on screen (control matching). For more details on the HCP task contrasts, see Barch, 2013 and the HCP 1200 Subjects data release reference manual (WU-Minn, 2017).

### Structural and functional predictors (prediction features)

We employed six types of cortical features as predictor modalities in our models: individualized resting-state functional connectivity measures, i.e. PCA (Hong, 2020) and ICA (Tavor, 2016; Ngo, 2021) component loadings, a measure of structural connectivity (Warrington, 2020), and structural measures (such as T1w/T2w, cortical thickness, curvature, and sulcal profile) serving as proxies for cortical microstructure and morphology, as well as three measures of cortical geometry, i.e. distances on the cortical surface (Wagstyl, 2015; Margulies, 2016, Oligschlager 2018, Wang, 2021) to reference landmarks, a PCA-decomposition thereof, and geometric eigenmodes (Pang et al. 2023), both derived from individual cortical surfaces. This section provides a detailed description of each feature type.

#### Resting-state functional connectivity PCA (RS-PCA) component loadings

For each subject, all four resting-state (RS) connectivity runs were z-scored (the standard deviation of the resulting values at different time points for a given vertex was thus ensured to equal 1) and concatenated along the time dimension (keeping the order of acquisitions with alternating phase encoding constant across sessions: S1LR, S1RL, S2LR, S2RL), resulting in a v x t matrix, with t=4800 at a TR of 0.7s. The timeseries of each cortical vertex was correlated with that of all other vertices, resulting in a (v x v) functional connectivity matrix C where c_i,j_ is the Pearson correlation coefficient between vertex i and j, and with v=29696 in the left brain hemisphere. The dimensionality of this matrix was then reduced to *d*=200 using Principal Component Analysis (PCA), resulting in i. an (v x d) matrix *W* that describes the basis vectors (components) of the subspace into which *C* was projected, and ii. a corresponding (v x d) matrix T_PCA_ that contains the loadings for each of these components at each vertex. Importantly, the components in *W* are orthogonal to each other, with each capturing a distinct (and progressively smaller) portion of the variance in *C*. We chose d=200 to match the number of ICA components (see next section). When possible, we also used d=200 for the other feature modalities. On average, the cumulative variance explained by these components was 84.7% (std=3.84%). Similar approaches have previously found wide applications in the field, e.g. in the form of functional gradients (Margulies, 2016, Huntenburg, 2018; Bernhardt, 2022; Hong, 2020). We then used Procrustes alignment — as implemented in the BrainSpace Python toolbox (Vos de Wael, 2020) — to align these loadings to a set of reference loadings. We derived the reference loadings by applying PCA to the group-level functional connectivity matrix provided by the HCP (Smith, 2014). This technique is used to ensure matching order, valence and rotation of the loadings across subjects. Procrustes alignment finds the transformation (allowing for rotations, translations, and scaling) for the individual loadings that minimizes the least square difference between the transformed loadings and the group loadings. The resulting group-aligned individual loadings were then used as model inputs.

#### Resting-state functional connectivity ICA (RS-ICA) component loadings

To enable comparison with previous studies (Tavor, 2016, Ngo, 2021), we also included Independent Component Analysis (ICA)-derived component loadings as a set of predictors. As in the case of PCA, we started with the z-scored and concatenated resting-state time series. We then correlated the time series of each cortical vertex with the individualized time series of each of the d=200 ICA group components (the highest granularity provided in the HCP release; derived through dual regression of the group-level ICA components onto the individual resting-state time series). We used the resulting matrix T_ICA_ of shape (v, d) as input to our model.

#### Distances on the cortical surface to landmarks

To derive our first measure of cortical geometry, we calculated the geodesic distance – that is the shortest distance on the cortical surface between two points – of each left hemispheric vertex to a range of landmarks. We computed these distances on the individual cortical surface (in the native space). Landmarks were defined as centroids of parcels (p=223) in a group-level parcellation (Lausanne 2008 scale 4, see Cammoun, 2012) spanning the entire left hemisphere. This parcellation subdivides the gyral-based Desikan-Killiany atlas (Desikan, 2006) into smaller parcels. We chose this parcellation, because of its medium granularity and similarly shaped parcels, to reduce any bias that could come from dissimilar parcel shapes or marked differences in the number of vertices in each parcel. In HCP, individual cortical surfaces are aligned to the same reference space (fsLR), so that individual vertices should approximately correspond to the same point on the cortex across subjects, which allows for usage of a group average based parcellation to parcellate the aligned native brain surfaces. Distances between vertices v (defining the space for the task maps) and parcel centroids p (=approximate landmark position) were measured on the individuals’ *native midthickness* cortical surfaces (halfway between pial and white-matter, rotationally aligned to fsLR32k space, and medial wall removed). This resulted in a single distance matrix of shape (v, p) per subject.

#### Cortical surface distance matrix PCA component loadings

As an alternative to the first parcellation-based approach, we computed the full vertex-to-vertex geodesic distance matrix, precluding any bias from the choice of a specific parcellation. Similarly to the RS-PCA approach, this full matrix was then reduced by PCA to 200 dimensions and aligned to a set of 200 reference loadings derived from the group average vertex-wise distance matrix. As the creation of a group-average surface likely leads to the loss of fine-grained spatial information due to blurring, we also alternatively aligned the components to distance components computed for reference subjects instead, which yielded comparable results.

#### Geometric eigenmodes of the cortical surface

As a third measure of geometry, we computed the first d=200 geometric eigenmodes (Laplace-Beltrami eigenmodes) on the individual cortical native midthickness meshes (excluding the medial wall) using the LaPy Python package using the lapy.solver.eigs function (in keeping with Pang 2023), resulting in a single matrix per subject of shape (v, d). These modes contain information about the distances between vertices, but also of other geometrical & topological features such as local curvature. Components hierarchically decompose the cortical mesh into increasingly fine-granular (periodic) maps (with the first modes typically differentiating between front-back, top-bottom, front-middle-back, and so on).

#### Structural connectivity blueprints

As a proxy for structural connectivity, we used a set of precomputed XTRACT connectivity blueprints (Warrington, 2020; available since FSL version 6.0.5). These blueprints describe maps of “cortical termination”, giving for each cortical vertex (v) an estimate how much each of the predefined k=41 fiber tracts contribute to the overall connectivity of that specific cortical location. We selected these blueprints because they offer a concise and comprehensive description of the brain’s structural connectivity, relate to established anatomical patterns (fiber tracts), and were demonstrated to capture inter-subject variability (Warrington, 2020). For each subject, a single matrix of shape (v, k) described its structural connectivity.

#### Cortical microstructure/morphology

As proxy for cortical microstructure and morphology, we used the maps provided as part of the HCP1200 data release in fsLR space at a resolution of 32k vertices for the left hemisphere. Morphological features were computed by the FreeSurfer recon-all routine. We focused on six features: the radiofrequency transmit field (B1+) bias-corrected (Glasser, 2022) ratio of signal intensity in T1-weighted and T2-weighted magnetic resonance images (T1w/T2w), as a measure of cortical microstructure (often taken as proxy for cortical myelin content, see Glasser & Van Essen, 2011), uncorrected and curvature-corrected cortical thickness, curvature, and sulc (FreeSurfer’s analog of sulcal depth). These features were organized into a single matrix of shape (v, 6) per subject, where *v* represents the number of cortical vertices.

### Prediction models

#### Vertex-wise prediction models

We used a single linear model per feature modality to do per-vertex predictions of task contrasts. That is, we input the feature values at a specific vertex in the brain and predict task contrast values at this specific vertex, whereby the number of features per vertex (=sample) ranges from 5 to 200, depending on the modality used as predictor. The full set of samples used for model fitting consists of all vertices of all training subjects. Parameters are thus shared across all subjects and all locations on the brain. The models thus may only take local information into consideration, and do not integrate any information over the spatial neighborhood. This contrasts with previous approaches, which often take features from multiple locations across the cortex as input (Langs, 2015; Tavor, 2016; Jones, 2017; Ngo 2022, Zheng 2022, Tik, 2023). For example, in these studies, 29k x 50 features per sample would be used to predict a full cortex-wide map at each prediction step, resulting in a much higher number of model parameters. Our method affords a much-increased sample size (29k samples per subject) and a marked reduction in model parameters (50 per task contrast), enabling estimation even on a small set of subjects, and easier interpretability.

Model parameters were estimated based on all left cortical samples from 203 subjects that formed the training dataset, resulting in a total of ∼6 million samples (203 subjects x 29k vertices), to predict ∼6 million activation values for each task. One model per task contrast was estimated using linear regression, resulting in a separate set of coefficients (one per input feature) for each task contrast. Features were z-scored across the spatial dimension. Assuming a simple linear model of Y’ = X × w, we derived the coefficients w of shape (d) that minimize the difference between the actual task contrasts Y and the predicted contrasts Y’, both of shape (n_subjs_ × v). The predictors X were of shape (n_subjs_ × v, d) being the concatenation of {T_subj1_, T_subj2_, …, T_subjN_} along vertices. Model fit is conducted using sklearn’s implementation of ordinary least squares Linear Regression as defined in the class ‘sklearn.linear_model.LinearRegression’. Evaluation of model performance is based on a held out set of 51 test subjects.

By virtue of using linear models, we can interpret the magnitude of the individual coefficients in w as a measure of how important each feature was for the prediction, as long as the model inputs (i.e. feature maps) are of the same scale (achieved by z-scoring them) and orthogonal to each other (as was the case for most of our feature modalities, including the RS-PCA and RS-ICA components). Due to their comparably small number, we decided not orthogonalize the microstructure/morphology maps to keep the direct correspondence with labels of the original maps. Hence small differences in the resulting coefficients should not be overemphasized. This approach contrasts with more complex models that might be able to capture nonlinear interactions between predictors, but these would not allow us to clearly delineate the contribution of each feature independently.

#### Per-parcel prediction models

To explore brain region specific associations between feature sets and task activations, instead of fitting a single brain-wide model per task, we fit one single linear model for each region (=parcel of a whole cortex parcellation) and task. For that we used the same parcellation as described before (Cammoun, 2012) containing 223 parcels. To fit each model, we first restricted both feature maps (predictors) and task contrast maps (prediction targets) to only the vertices that belong to a given parcel (resulting in a training set of e.g. 203 subjects * 120 vertices ∼ 24360 samples, if the parcel contained 120 vertices). In this case, all scores were computed on the restricted set of vertices in the test subjects.

### Measures of model performance

To quantify model performance, we used three main measures (Figure 1E-F): Overall **prediction accuracy** defined as the correlation between task maps was computed as the average correlation of the true task maps and the model predictions across individuals in the test set (Eq.1). This yielded one value per test subject which can then be averaged across subjects. Task maps were one-dimensional arrays of length v (with a mean of zero and a standard deviation of one), describing the z-scored activations at each cortical vertex. Given that we can achieve a high accuracy score close to group-average baseline (merely “predicting” the same group-average task map for each individual), we furthermore assessed the **prediction discriminability** (similar to Zheng et al 2022) which was defined as (i) the average correlation between the predicted and the actual map of each individual from which we subtracted (ii) the average correlation between an individuals’ predicted maps and and all other individuals’ maps in the test set (Eq. 2). This yielded a single value per test subject. This measure allowed us to assess the model’s ability to capture individual differences.

As a third measure, we computed the **vertex-wise accuracy** as the correlation between the series of predicted task contrast values at a given vertex for all subjects (a vector containing one value for each subject) and the series of corresponding true values (Eq. 3), yielding one value for each vertex (across all test subjects). This vertex-wise accuracy allows us to assess in which parts of the brain the model accurately predicts individual differences.

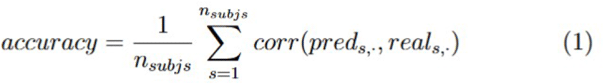

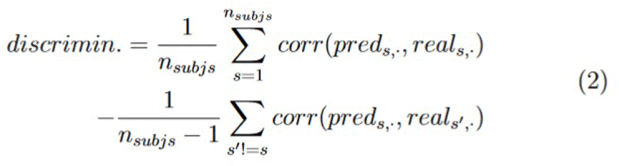

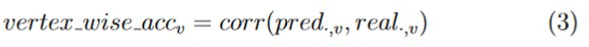

#### Prediction baselines

To put these scores into context, we calculated two types of baseline scores on the test-retest subset of HCP. First, we calculated the **group-average prediction baseline** for the prediction accuracy scores by replacing each of the predicted task activation maps by the corresponding group average task activation map (task activation maps averaged across individuals) and then calculating the scores. This corresponds to the baseline model accuracy or discriminability if no individual predictions were done, but instead the same group average contrast map would be predicted for each subject. By design both the prediction discriminability and the vertex-wise accuracy scores are zero when always predicting the same task map, so we did not report them separately. Furthermore, we computed the **test-retest baseline** for all the three scores by basing their calculation on the test and retest contrast maps (available for a subset of HCP subjects), instead of real and predicted contrast maps.

### Relation of resting-state components with other cortical maps

To understand what kind of functional and biological characteristics may be related to the resting-state components, we correlated each of the group components with 101 group-level cortical reference maps. This adds context to the interpretation for why a specific component could be important for the emergence of a function, and may provide a link to features that can not or have not yet been assessed at the individual level.

These reference maps encompass the 7 and 17 yeo networks (Yeo, 2011), the alternative 12 Cole-Anticevic networks (Ji, 2019), three robust gene expression components (Dear, 2023) along with 12 further components compiled by the same author (pertaining e.g. to T1/T2 and externopyramidisation), the 10 principle functional connectivity gradients (Margulies, 2016) downloaded from neuromaps, 24 cell type maps (Zhang, 2023; Jorstad, 2023) and 16 receptor maps (Hansen, 2022). In some cases (i.e. the cell type maps), data was only existing on the regional (=parcellated) level, e.g. due to limitations in the method. In those cases these maps were upsampled to fsLR29k (without interpolation) prior to correlation and spin tests.

We report the most strongly correlated reference maps (by magnitude of correlation) for the top 4 components predictive for each task contrast map, along with their correlation coefficient and p-value. The latter were derived using a spatial spin-test (Alexander-Bloch, 2018) with 5000 permutations, as implemented by the BrainSpace toolbox. This approach comes with limitations (Leech, 2024), but was chosen for its simplicity and computation efficiency in face of the high number of correlations performed. All the reported p-values are false discovery rate (FDR)-adjusted using the Benjamini-Hochberg method (Benjamini & Hochberg, 1995) across tests performed within each category of reference maps. Furthermore, our interpretation of these correlation focuses on the differences in correlation strength and not the significance of the correlation per-se.

### Validation of our findings across hemispheres and datasets

To ensure the robustness of our findings, we repeated our analysis in the right hemisphere. A separate model was fitted using the corresponding data from the right hemisphere, following the same methodology outlined for the left hemisphere. This allowed us to compare the task-specific component patterns between hemispheres, ensuring that our results were consistent and not hemisphere-specific. Even though there might be asymmetries in the task activation patterns for specific tasks that show greater engagement of one hemisphere, it should still be possible to predict variations in activity patterns in the non-dominant hemisphere.

Second, we conducted an external validation using the HCP-style data from the Courtois project on Neural Modelling (Courtois NeuroMod). We will refer to it as the CNM dataset throughout the manuscript. The CNM dataset provides preprocessed resting-state and structural measures in the appropriate format. To derive task contrast maps, we used the preprocessed task runs (n=4, selecting every second task run) and applied the task contrast pipeline implemented in Nilearn. We focused on the most replicable task contrasts (as described previously) that could be computed based on the event annotations provided with these task runs. Additionally, we assessed test-retest baseline scores using the remaining four task runs. For the computation of task contrasts, we fitted a first-level GLM to each task run using Glover’s model for the hemodynamic response function along with a time derivative (“Glover+time”), a cosine drift model, and a high-pass filter frequency of 0.005. Fixed-effect t-statistic task contrasts were then computed based on these first-level results.

## Results

Building upon previous research focused on single-modal predictions (Langs, 2015; Tavor, 2016; Jones, 2017; Tobyne, 2019; Ngo, 2022; Zheng, 2022; Tik, 2023), we employed a comprehensive range of feature modalities to elucidate the intricate relationship between cortical features and functional activations. This involved fitting predictive models based on individualized resting-state PCA and ICA component loadings, structural connectivity, and structural measures (including intracortical myelin, cortical thickness, curvature, sulcal profile), and measures of cortical geometry.

In contrast to common approaches that treat each participant as a single data point, our lightweight prediction model considers every cortical vertex as an individual data point - similar to models employed by Saygin (2012) and Tobyne (2018). By virtue of this, it requires only a few participants for model fitting and offers enhanced interpretability. The small number of weights directly linked to individual features readily enables the analysis of the contributions of specific features within each modality to the prediction success and their relevance in cortical functional organization.

In the presentation of the results, we focus on the task contrasts with the highest test-retest discriminability for each of the seven tasks assessed for HCP. This includes the following contrasts: Language Story-Math, Motor Cue-Avg, Emotion Faces-Shapes, Working Memory 2Back-0Back, Social Theory of Mind-Random, Relational Rel-Match and Gambling Reward-Punish. For brevity, we refer to them only by their task name.

### Resting-state derived measures best predict individual task activation locations

To identify which feature modalities most effectively capture differences in functional activations, we began by fitting separate linear models for each modality using a training set of 203 participants. For each model, we quantified overall prediction accuracy by calculating the average correlation between the actual and predicted maps. Additionally, we assessed discriminability — the difference of the average correlation of predicted and actual subjects’ task maps, minus the average correlation of predicted and any other subjects’ target map — to determine how well each modality captured individual differences in functional activations. Moreover, we computed vertex-wise accuracy — the correlation of real and predicted values across subjects at each cortical vertex — to identify the parts of the cortex in which the models are performing the best. These metrics were calculated based on a held-out test set of 51 participants, allowing us to compare the predictive power of each modality. We ground these metrics by two baselines: a test-retest baseline — providing information of how stable individual task maps are across two separate sessions, according to a given measure — and a group-average baseline, indicating which score could be obtained if always the same group average map was predicted.

Individual resting-state functional connectivity PCA loadings (RS-PCA) yielded the highest accuracy (r_language_=.688, r_motor_=.594, r_relational_=.318, r_gambling_=.102) and discriminability (Δr_language_=.196, Δr_motor_=.134, Δr_relational_=.094, Δr_gambling_=.033) across tasks, tracking test-retest baseline scores for both accuracy (r_language_=.704, r_motor_=.6, r_relational_=.195, r_gambling_=.048) and discriminability (Δr_language_=.268, Δr_motor_=.225, Δr_relational_=.11, Δr_gambling_=.035). The vertex-wise accuracy pattern for the RS-PCA-based model mirrors closely the test-retest baseline, suggesting good predictivity for task contrast differences across the cortex (Supplementary Figure 2). Similarly individual ICA loadings were highly predictive of the task maps measured by both accuracy (r_language_=.689, r_motor_=.583, r_relational_=.288, r_gambling_=.092) and discriminability (Δr_language_=.182, Δr_motor_=.121, Δr_relational_=.07, Δr_gambling_=.032) close to those of RS-PCA.

Contrary to our initial hypothesis, the models based on the remaining feature modalities – including distance measures, cortex structure maps, geometric eigenmodes and anatomical connectivity yielded discriminability scores close to zero (e.g. Δr_language_=.001, Δr_motor_=-.004 for distances), despite good accuracies (r_language_=.471, r_motor_=.513 for distances), on par with group average baselines (r_language_=.665, r_motor_=-.624),. This indicates that group average maps can be approximated (reflected in the accuracy score), but no individual differences can be accounted for by any of these modalities (reflected in discriminability). These results are summarized in Figure 2 and example predictions for some of the contrasts are shown in Figure 3. The scores for the remaining tasks can be found in Supplementary Figure 1. Per-parcel models (Supplementary Figure 4) largely confirm these findings.

**Figure 2.**
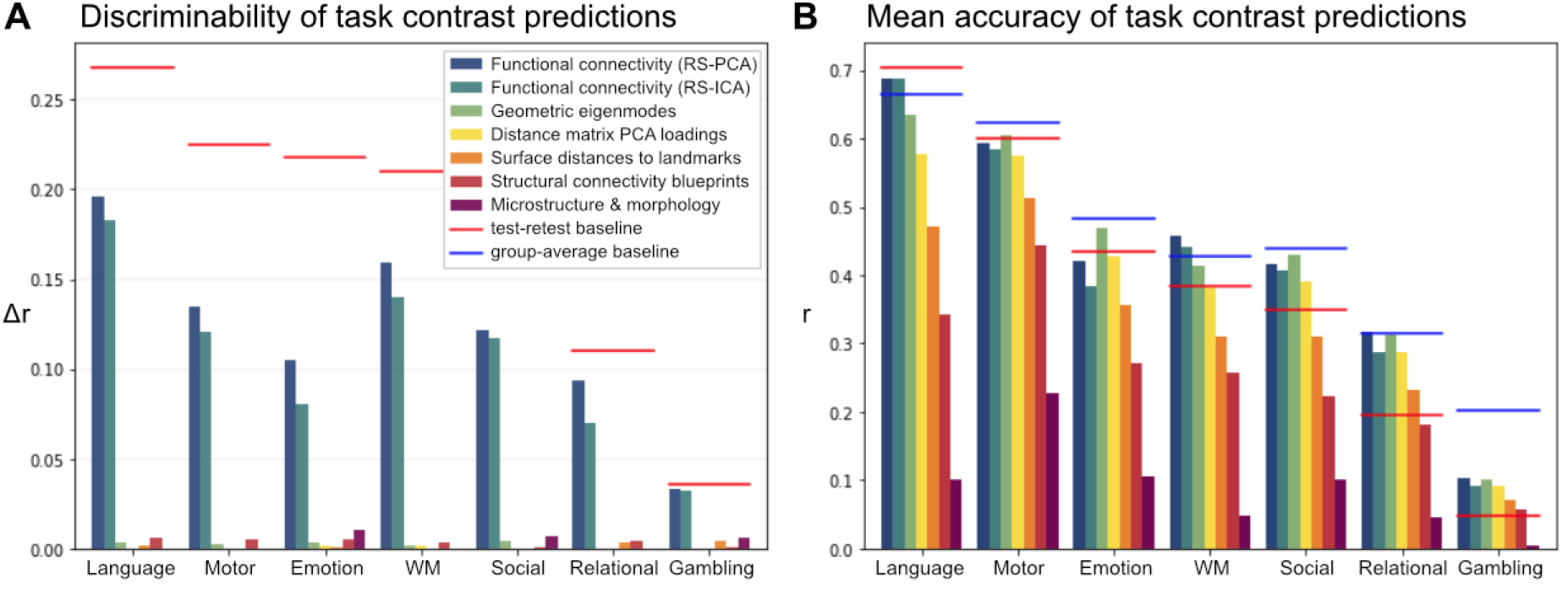
**A** Discriminability of task contrast maps, quantifying how well individual differences are captured. **B** Accuracy of predicted task contrast maps. This score quantifies how well the overall group-level task distribution can be captured, but does not differentiate between capturing the group-level distribution or individual differences. Scores are shown for the task contrast with the highest test-retest discriminability for each of the seven tasks assessed for HCP. This includes the following contrasts: Language Story-Math, Motor Cue-Avg, Emotion Faces-Shapes, Working Memory 2Back-0Back, Social Theory of Mind-Random, Relational Rel-Match and Gambling Reward-Punish. Only the task names are shown in the figure for brevity. For a more complete description of the tasks see Barch (2013). Horizontal lines indicate both the test retest baseline (aimed at measuring the stability of task contrast maps in individuals across sessions, providing a noise ceiling for each task contrast for both discriminability and accuracy) and the group-average baseline (i.e. the scores that would be obtained if always only the group-average task map was was predicted, and by construction equaling zero in the case of discriminability). The apparent overlap in both baselines in panel B evidences that the mean accuracy - in contrast to the discriminability score - does not allow to distinguish whether or not the models are capturing individual differences.

**Figure 3.**
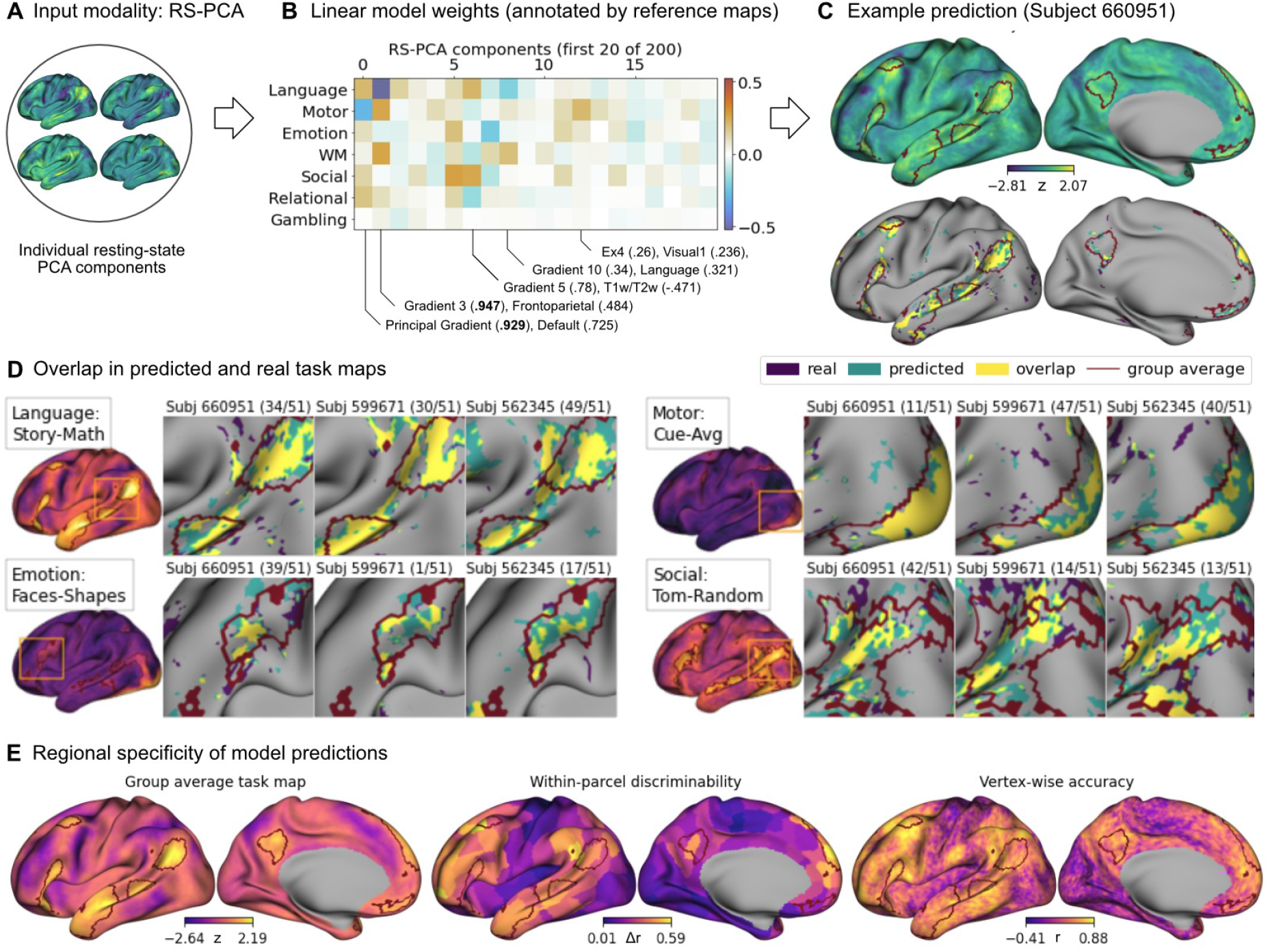
Resting-State PCA-based prediction of full task maps using a linear model. **A** The model takes group-reference aligned individual resting-state PCA component loadings as input. **B** The coefficients for these loadings are estimated in a task-specific manner. Individual component loadings are aligned to group-level component loadings prior to model fit, ensuring that they match across subjects. Most predictive components for both the language and motor task are annotated (below the plot) by the highest correlating reference maps at the group-level, highlighting possible relationships to functional gradients (Gradients 3, 5 and 10), functional networks (Visual, Language, Default and Frontoparietal), cell types (Excitatory Type EX4) and microstructure (T1w/T2w). **C** The upper row contains the predicted map for a sample contrast (Language Story-Math) for a sample subject with intermediate individual discriminability based on the model (rank 34 out of 51 subjects, with lower rank indicating higher discriminability). The lower row depicts the thresholded (above the 90th percentile) real (purple) and predicted (green) maps for the same subject, highlighting the overlap between the two in yellow. **D** Individual differences in task contrasts are accurately reproduced by the model. The thresholded real and predicted task maps (similar to panel C) are displayed for three sample subjects across 4 different task contrasts (right) with cutouts focusing on regions that show high activation in the corresponding group-average contrast map (left). The labels indicate subject number and rank. **E** From left to right: The group average map for the Language Story-Math contrast as reference. Within-parcel discriminability scores are computed based on the real and predicted values by the full model for only the vertices that belong to a given parcel (Supplementary Figure 3). This visually mirrors the group-average task map, indicating that the model predicts individual differences best in task-specific regions. This is confirmed also by the vertex-wise accuracy scores for the model that closely mirrors test-retest vertex-wise accuracy (Supplementary Figure 2).

### Task-related functional activations are associated with specific resting-state connectivity and patterns

To understand which specific RS-PCA component loadings were predictive of task contrast differences, we looked at the weights of the linear model. By prior z-scoring the input data along the spatial dimension we ensured that the betas for different features/components should be within a comparable range. This yields a set of most strongly weighted components for each task contrast (Figure 3B; Supplementary Figure 5). Predictive components were often shared among multiple tasks (e.g. c0 and c1), and the components that explained more variance in functional connectivity also had higher absolute betas. Some components were anticorrelated across tasks, possibly reflecting the exclusive or opposing nature of these tasks (e.g. β_c0_=.205 in the language and β_c0_=-.288 in motor contrast). Other components were more task specific, such as c12 for the motor contrast (β_c12_=.206). Interestingly, task-over-baseline contrast maps, such as story, math, cue, or avg were predicted by a very similar set of components (most prominently c1,c3 and c5), possibly explaining individual differences in activity related to general task engagement which likely overshadowed smaller task-specific activations.

**Figure 4.**
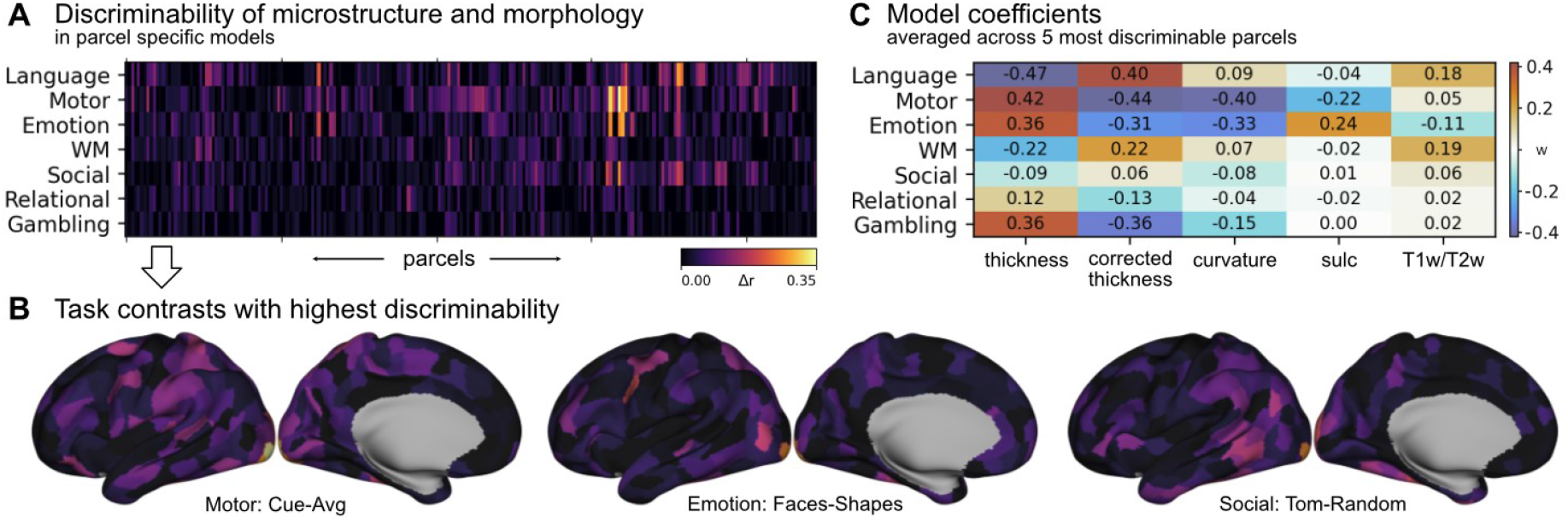
Microstructure and morphology based prediction using per-parcel models. **A** Discriminability for each of the per-parcel models for the seven main task contrasts. **B** Discriminability scores from A plotted on the cortical surface for three task contrasts, showing that highest discriminability can be found in occipital regions. **C** Weights for the linear model predicting task maps, averaged across the five most discriminable per-parcel models, highlighting cortical thickness and curvature as predictive features.

**Figure 5.**
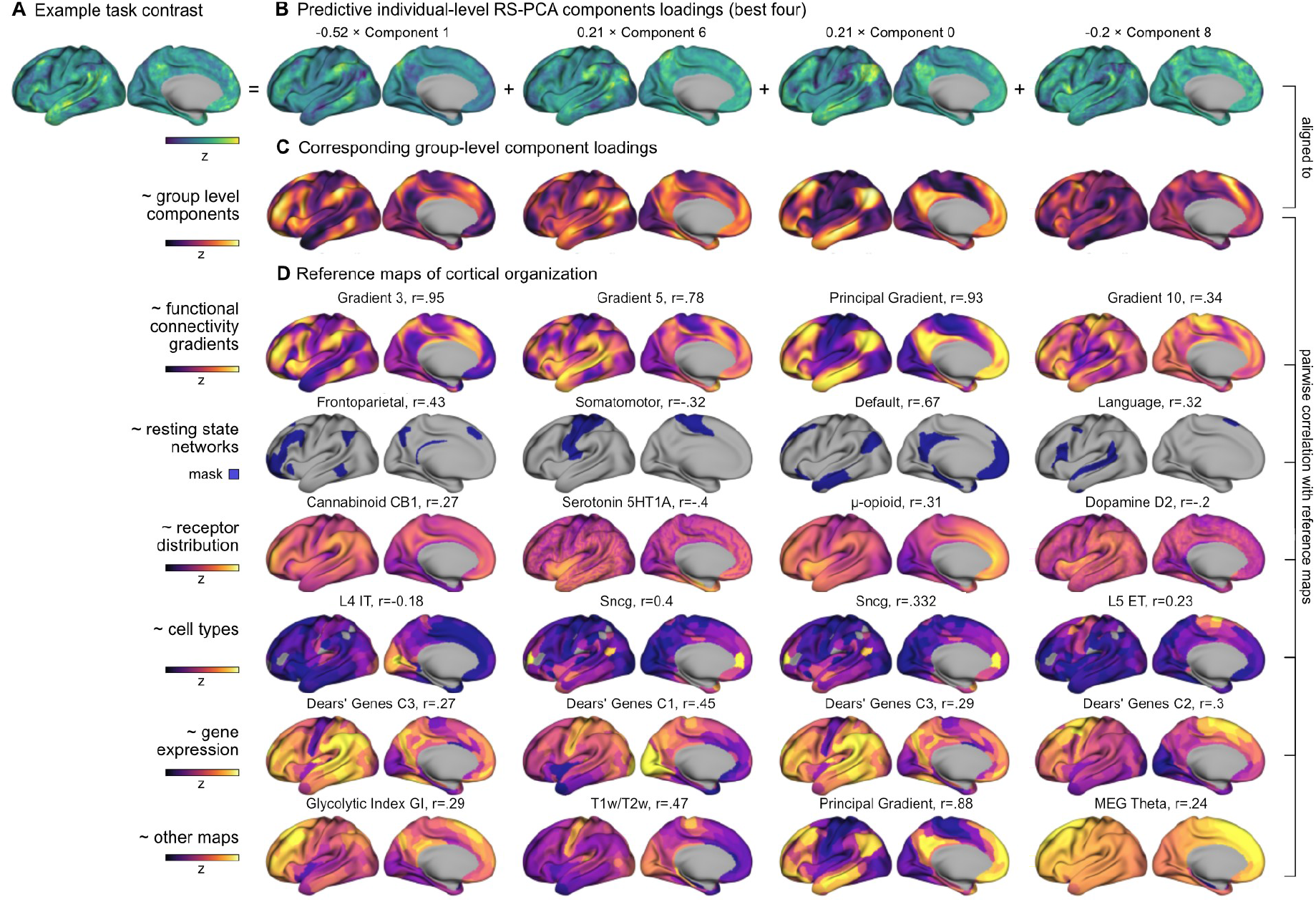
Task predictive resting-state components can be related to different reference maps. **A** The predicted contrast map (on the example of Language Story-Math). **B** This task contrast map is composed as the sum of individual-level resting-state components weighted by the task specific model coefficients. **C** For each of the individual components we show the corresponding group-level component to which these were aligned prior to model fit. **D** Each of these group-level components can be linked to reference maps. We highlight the most strongly correlated maps (based on Pearson’s r) for each type of reference (resting-state functional connectivity gradients, functional networks, receptor distribution, cell types, gene expression, and other maps derived from neuromaps). These constitute candidate constraints for functional properties that are relevant across tasks.

### Cortical thickness is predictive of function within task-specific subregions of the cortex

In addition to brain-wide models, we also ran a set of per-parcel models for each of the feature modalities. This allows capturing of local relationships between features and functional activations that may not be present throughout the cortex. We find that in addition to RS-PCA, cortical structure (i.e. cortical thickness, curvature & myelin content, assessed using freesurfer) was also predictive of differences in task activation, especially in the motor task (Figure 4a-b, all feature modalities are shown in Supplementary Figure 4). This seems reminiscent of the slightly elevated discriminability scores in the full brain models for the emotion (Δr_faces-shapes_=.01) and social (Δr_tom-random_=.007) task contrasts. The per-parcel predictions are mainly driven by cortical morphology, as evidenced by the feature coefficients averaged across the 5 per-parcel models with the highest discriminability (Figure 4C). For example, the most predictive features in the motor task are uncorrected (β=.42) and corrected cortical thickness (β=-.44) and curvature (β=-.4). Plotting the discriminability scores of the per-parcel models on the cortical surface revealed that cortical microstructure/morphology was mainly predictive of activations in the occipital cortex (Figure 4B).

### Task predictive resting-state components can be related to group-level functional networks, gene co-expression patterns, receptor and cell type distribution

To render the specific resting-state components (c0, c1, … c200) that we found predictive of task activations more interpretable and to link them to possible neurobiological substrates, we characterize each resting-state component through correlation analysis (Figure 5, Supplementary Figure 5). We focused this analysis on the resting-state components, because unlike the microstructural predictors such as cortical thickness that are inherently linked to the brain’s physical properties, these components do not have an obvious and clear connection to underlying neurobiology. For that, we correlated each of the group-level components with a set of group-level cortical maps representing functional networks, gene co-expression patterns, resting-state functional connectivity gradients, receptor and cell type distributions. We find that, for example c0, whose loadings were relevant for the prediction of multiple task contrasts, was most strongly correlated with the first resting-state gradient (r=.929) and the default mode network (r=.725), reflecting the similar derivation of our RS-PCA components and typical resting-state gradients. Similarly, c1 - another component relevant in predicting multiple task contrasts - was most highly correlated with gradient 3 (r=.947) and the frontoparietal network (r=.484). Looking at two exemplary main task contrasts and their non-shared most predictive components, we found that in the language contrast, c6 (β=-.21) may be related to the T1/T2 ratio (r=-.471), the first component of gene expression (r=-.45), and a range of cell and receptor types, e.g. synuclein gamma -like interneurons (SNCG, r=.402), parvalbumin-expressing interneurons (PVALB, r=-.394) and serotonin receptor 1a (5HT1a, r=.402), and c8 (β=-.201) might be related to the language network (r=.321) and the second gene expression component (r=.3). Also the WM-contrast had high betas for c8 (β=.187), possibly linking it to the language network as well. The motor contrast uniquely draws on c12 (β=.206), which correlates with the layer 4 intratelencephalic-projecting excitatory neurons (L4 IT, r=.251) and the first visual network (r=.236), mirroring this contrasts’ selectivity for visual areas. The emotion contrast had high betas for c7 (β=-.223), correlating with gradients 6 (r=.533), 7 (r=.403) and 10 (r=.381), dopamine transporter DAT (r=.454) and anticorrelated to serotonin transporter 5HTT (r=-.399), and for c5 (β=.188) which may be associated with gradient 2 (r=.69) and is anticorrelated to the somatomotor network (r=-.577). A full listing for all the task contrasts can be found in Supplementary Table 2 and Supplementary File A.

### The same resting-state components predict individual differences in both hemispheres

In the previous sections, our description of the results primarily focused on the left hemisphere to explore the predictive relationship between resting-state components and individual differences in task activations. To ensure the robustness of our findings and assess potential lateralization effects, we validated our findings in the right hemisphere. Overall we found a similar pattern for accuracy and discriminability across left (e.g. r_language_=.688, r_motor_=.594, r_relational_=.318, r_gambling_=.102, Δr_language_=.196, Δr_motor_=.134, Δr_relational_=.094, Δr_gambling_=.033) and right hemispheres (r_language_=.694, r_motor_=.534, r_relational_=.317, r_gambling_=.119, Δr_language_=.21, Δr_motor_=.149, Δr_relational_=.074, Δr_gambling_=.037). We further sought to verify whether the same resting-state components that predict individual differences in the left hemisphere are also predictive in the right hemisphere. We found an overall similar pattern — e.g. for the language (β_left_c1_=-0.52, β_right_c1_=-0.55) and motor (β_left_c1_=0.25, β_right_c1_=0.19) contrasts. The correspondence becomes more robust after additional Procrustes alignment of the coefficients from the right hemisphere to those of the left to correct for differences that stem from computing the group-level reference components separately for each hemisphere. The resulting aligned coefficients correspond across hemispheres — e.g. the most predictive components for the language contrast in the left were c1 (β=-0.523), c6 (β=0.214), c0 (β=0.206), and in the right c1 (β=-0.506), c6 (β=0.22) and c8 (β=-0.214). For the motor contrast, the most predictive components were c0 (β=-0.288), c1 (β=0.254) and c12 (β=0.206) in the left, and c12 (β=0.245), c0 (β=-0.236) and c1 (β=0.211) in the right. The full overview of the first 30 components across hemispheres can be found in Supplementary Figure 6. This confirms that the same resting-state components were predictive of individual task activation differences in both hemispheres, highlighting their role in brain function independent of the hemisphere.

### The resting-state component based model generalizes across different datasets

Until now, our analysis focused solely on subjects from a single dataset (HCP). To substantiate our results, we probed the models’ ability to generalize to independently acquired datasets. To this end, we applied the model, originally fit on the HCP dataset, to a new dataset comprising three subjects from the Courtois NeuroMod project, referred to as CNM dataset in the remainder of the section. For that, we first manually calculated the task contrasts, limiting ourselves to those with the highest test-retest reliability in the HCP dataset that were readily computable from the available data, thus excluding the motor contrast. Notably, we observed a similar prediction pattern albeit with slightly smaller scores in CNM (r_language_=.688, r_relational_=.318, r_gambling_=.102, Δr_language_=.196, Δr_relational_=.094, Δr_gambling_=.033) as seen in HCP (r_language_=.426, r_relational_=.232, r_gambling_=-.013, Δr_language_=.164, Δr_relational_=.052, Δr_gambling_=-.011). We found the strongest deviation for the social task contrast in CNM (r_social_=.009, Δr_social_=-0.047) compared to HCP (r_social_=.417, Δr_social_=.122), which could be due to differences in how the task contrasts were derived. Given the very low number of test subjects (n=3), these results should only be seen as indicative. The full results can be found in Supplementary Figure 7.

To further validate our findings, we also assessed whether a model fit exclusively on the new dataset could identify similar components predictive of individual differences. Since we only had access to the data of three subjects, we fit our model on only a single subject, to allow for the computation of discriminability scores (which requires n>1) on the remaining two subjects. This again yielded comparable patterns in CNM (r_language_=.493, r_relational_=.06, r_gambling_=-.012, Δr_language_=.235, Δr_relational_=-.072, Δr_gambling_=-.004, also shown in Supplementary Figure 7) to the results on HCP, albeit unable to capture the relational task contrast (most likely due to only being fit on a single subject). In addition, our analysis revealed that similar resting-state PCA components were indeed predictive in this new dataset as well, at least for the well discriminable task contrasts, with e.g. the most predictive components for the language contrast in the HCP model being c1 (β=-.523), c6 (β=.214), c0 (β=.206), and most predictive aligned coefficients in the single subject CNM model being c1 (β=-2.222), c0 (β=0.964), c2 (β=.757), or for the emotion contrast in the HCP model being c7 (β=-.223), c5 (β=.188), c11 (β=.125), and in CNM being c7 (β=-1.187), c23 (β=.713) and c5 (β=.622). The change in scale is likely a result of fitting the model on only a single subject. The full results can be found in Supplementary Figure 8. This not only corroborates the features identified by our initial model but also demonstrates that such a model can effectively be fitted on very limited data. This capability is particularly advantageous for potential practical applications where data availability may be constrained.

## Discussion

In this study, we present a versatile analytic framework for exploring the relationship between brain structure and function. By evaluating a diverse range of feature modalities, we found that functional connectivity as well as cortical microstructure/morphology are closely linked to layout of functional specialization at the individual level. We specifically revealed a significant predictive relationship between individual spatial loadings of resting-state PCA components and task activations, with both the model and the patterns of the predictive components generalizing across hemispheres and to independent datasets. The association between cortical microstructure/morphology and functional activations was specific to the occipital cortex, highlighting their nuanced interplay with cortical types.

While lower-order resting-state functional connectivity PCA components were found to be informative of the overall distribution of functional activations across tasks, our study emphasizes the relevance of higher-order components in predicting task-specific functional distributions. Much prior research has focused on the first few components— often referred to as functional gradients —and their links to neurobiological features such as cortical microstructure, semantics, processing timescales, gene expression (Huntenburg, 2018; Bernhardt, 2022), and developmental plasticity (Sydnor, 2023). Fewer studies have investigated the full range of components (Hong, 2020). Our analysis links both lower and higher order resting-state components with functional activations and characterizes their possible neurobiological basis. For instance, we observe that the component predictive of individual spatial distribution in the language contrast may exhibit associations with both the resting-state language network and the second gene expression gradient identified by Dear et al. (2023). Similarly, we find that a specific class of excitatory neurons (L4 IT) characterizes the resting-state component that coincides with the primary visual network and predicts individual functional activations elicited by visual cues.

These findings corroborate previous studies which reported a relationship between resting-state fluctuations and task-specific activations (Fox, 2005; Cole, 2016). Additionally, they extend our understanding by demonstrating the network-specific nature of task activations, possibly grounded in underlying neurobiological features such as gene expression and cell types. Interestingly, a previous study has found that the location of the visual word form area can be predicted from structural connectivity before its functional selectivity is established (Saygin, 2016), showcasing that neurobiological features may indeed precede and likely shape functional selectivity. Similarly, functional connectivity networks, being grounded in intrinsic neurobiological features, could delineate or even precede the formation of functionally specialized regions in the cerebral cortex.

We also found that individual differences in task-over-baseline contrast maps are predicted by a similar set of resting-state functional connectivity components across contrasts. These components likely serve to explain individual differences in activations related to general task engagement. Such activations may dominate the signal and possibly overshadow smaller task-specific activations. This may be indicative of an overall larger prominence of intrinsic brain activity underlying major functional networks, compared to task specific activity.

Given the usefulness of the connectivity components in characterizing individual differences in functional activations across tasks, they may potentially also be useful for functional fingerprinting (Finn, 2015). Interestingly, one of the most predictive components of individual functional activations across tasks in our model closely mirrors the frontoparietal network. The same network had been previously identified as most distinctive across individuals (Finn, 2015) evidencing its ability to capture interindividual differences, consistent with our findings.

While we focused our analysis on the RS-PCA component loadings due to their inherent ordering and similarity to functional gradients - which rendered their interpretation more straightforward - we also found that RS-ICA components were nearly equally predictive. This may be because both types of components captured similar functional networks. Further, in line with previous studies, we found that different task contrasts were differentially predictable (with the language contrast being predicted best, and the gambling contrast least). This closely corresponds to the stability of these contrasts in individuals across sessions, as quantified with the test-retest baselines. Another observation in our study was that even contrasts that typically show lateralization (such as the language contrast) are equally well predictable in both hemispheres. This does not mean that the contrast has the same strength or extent in both hemispheres, but rather, that whatever can be predicted of the task contrast in a given hemisphere - however small it may be - is similarly well predicted in the left and the right hemisphere.

As part of our comprehensive analysis, we investigated cortical microstructure/morphology as a feature modality for predicting individual functional activations in specific tasks. While its predictive power was observed to be modest and was mainly restricted to the occipital cortex, its emergence as a spatially circumscribed predictive factor underscores the intricate relationship between cortical architecture and functional specialization. This finding aligns with previous research indicating an increased divergence between microstructure and function in higher-order regions (Huntenburg, 2017, Paquola, 2019).

In contrast to our initial hypothesis, we did not find evidence for a role of individual cortical geometry in predicting individual functional activations, despite prior studies indicating an association between cortical geometry and function at the group-level (Margulies, 2016; Pang, 2023). This broad association may stem from developmental processes (Chandra, 2024) guided by growth and patterning centers (Cadwell, 2019; Molnár & Kwan, 2024), which shape cortical neural differentiation in a spatial manner. However, the fine-grained functional layout may be determined by a myriad of other factors which may explain the absence of a relationship at the level of individual differences. Nonetheless, it is important to note that such relationships have been observed by others (Wang et al., 2023), which might be attributable to methodological differences.

Similarly, we did not find any association between structural connectivity and individual task activations, contrary to previous studies (Saygin, 2012, Saygin, 2016, Wu, 2018). This may be a result of differences in the processing of the diffusion-weighted MRI (DWI) data or their usage of a parcellation which may be more suitable for diffusion data, given its probabilistic nature and comparably low spatial precision. Given the relative sparsity of studies investigating the prediction of function based on geometry and structural connectivity, further investigation may be needed to detail their relationship to functional activations.

A shortcoming of our vertex-wise linear modeling approach is that it assumes that associations between task activations and prediction features are stable/fixed across the entire cortex. More complex models could increase the predictive accuracy, possibly by accounting for non-linear relationship between features and task activations. While our study benefits from the individual-level characterization of resting-state components and task activations, it is important to acknowledge that the linking to reference maps was constrained to group-level due to data availability. Whilst these group-level maps already provided valuable context, future research could gain deeper insights by incorporating individual-level assessments of additional cortical features, such as those obtained from magnetic resonance spectroscopy, and directly relating them to functional data. Such an approach would enable more precise testing of the associations suggested by our analysis, allowing for a more nuanced understanding of the relationship between cortical structure and function. Furthermore, reference maps differed in granularity (i.e. some were parcellated) and in their preprocessing (e.g. projection from volumetric to surface space), limiting the absolute comparison of correlation coefficients to within a given category of reference maps.

Our findings on the predictive capabilities of both resting-state components and microstructure/morphology may also be of clinical relevance. For instance, the modalities or features that prove most effective in predicting interindividual differences in function may also hold promise as potential disease biomarkers. Our approach to identifying and interpreting most predictive features in our models aligns with strategies used to identify biomarkers for conditions such as psychopathology (Chen, 2020; Park, 2020; Yin, 2023), aging (Hoffmann, 2023), and cognition (Finn, 2015; Hong, 2020; Ouyang, 2020; Xia, 2023). By linking these predictors to anatomical features we can enhance their interpretability and potentially better pinpoint their link to disease (Bazinet, 2023). Further the ability to forecast individual functional activations based on specific feature modalities may enhance presurgical mapping, offering potential improvements in feasibility, speed, and accuracy.

Beyond clinical applications, a better understanding of which features and biological characteristics contribute to cognitive functions could also motivate the inclusion of similar neurobiological and anatomical principles in artificial systems that aim at reproducing human-like functions. Expanding this predictive framework to include additional modalities and datasets in the future could further elucidate what features drive areal specialization in the cerebral cortex for complex functions.

## Supporting information

Supplementary File A - RS component annotations

## Code and data availability

We publish the code for our models, data processing and analysis on github (https://github.com/rscgh/individual_function_predictions) and encourage other research groups to validate our findings, and to transfer these methods to novel predictor modalities, model types and datasets.

## Acknowledgements

This research was funded by the European Research Council (ERC) under the European Union’s Horizon 2020 research and innovation programme (Grant agreement No. 866533) awarded to D.S.M. R.S. was funded by the University of Leipzig, the German Federal Ministry of Education and Research (BMBF) and the Max Planck Society. Data were provided in part by the Human Connectome Project, WU-Minn Consortium (Principal Investigators: David Van Essen and Kamil Ugurbil; 1U54MH091657) funded by the 16 NIH Institutes and Centers that support the NIH Blueprint for Neuroscience Research; and by the McDonnell Center for Systems Neuroscience at Washington University. Further data were provided by the Courtois project on neural modelling, which was made possible by a generous donation from the Courtois foundation, administered by the Fondation Institut Gériatrie Montréal at CIUSSS du Centre-Sud-de-l’île-de-Montréal and University of Montreal. The Courtois NeuroMod team is based at “Centre de Recherche de l’Institut Universitaire de Gériatrie de Montréal”, with several other institutions involved. See the cneuromod documentation for an up-to-date list of contributors (https://docs.cneuromod.ca). We further thank Pierre-Louis Bazin, Demian Wassermann and the team of Pierre Bellec (SIMEXP lab) for discussions on the project, and Shaun Warrington from the Computational Neuroimaging Laboratory (CoNI Lab), University of Nottingham for providing us with the connectivity blueprints.

## Supplementary Materials

**Supplementary Table 1.** Test-retest discriminability by task contrast.

| Contrasts | $\Delta r$ | Contrasts (continued) | $\Delta r$ |
| --- | --- | --- | --- |
| **LANGUAGE_STORY-MATH** | **0.268** | WM_OBK_PLACE | 0.183 |
| WM_2BK | 0.260 | MOTOR_RF | 0.182 |
| SOCIAL_TOM | 0.256 | WM_OBK_TOOL | 0.180 |
| WM_FACE | 0.251 | EMOTION_FACES | 0.177 |
| SOCIAL_RANDOM | 0.247 | MOTOR_LH | 0.177 |
| GAMBLING_REWARD | 0.243 | LANGUAGE_STORY | 0.174 |
| GAMBLING_PUNISH | 0.243 | **SOCIAL_TOM-RANDOM** | **0.169** |
| WM_2BK_FACE | 0.240 | EMOTION_SHAPES | 0.169 |
| WM_BODY | 0.229 | MOTOR_RH | 0.161 |
| WM_OBK | 0.227 | MOTOR_T | 0.145 |
| RELATIONAL_REL | 0.225 | WM_FACE-AVG | 0.138 |
| **MOTOR_CUE-AVG** | **0.225** | WM_PLACE-AVG | 0.126 |
| RELATIONAL_MATCH | 0.222 | MOTOR_T-AVG | 0.122 |
| WM_TOOL | 0.221 | WM_BODY-AVG | 0.116 |
| MOTOR_AVG | 0.218 | **RELATIONAL_REL-MATCH** | **0.110** |
| **EMOTION_FACES-SHAPES** | **0.218** | MOTOR_LH-AVG | 0.096 |
| WM_PLACE | 0.213 | MOTOR_RH-AVG | 0.094 |
| WM_2BK_BODY | 0.212 | MOTOR_LF-AVG | 0.087 |
| WM_2BK_TOOL | 0.211 | WM_TOOL-AVG | 0.083 |
| **WM_2BK-OBK** | **0.210** | MOTOR_RF-AVG | 0.077 |
| MOTOR_CUE | 0.204 | **GAMBLING_REWARD-PUNISH** | **0.035** |
| WM_2BK_PLACE | 0.201 |  |  |
| LANGUAGE_MATH | 0.194 |  |  |
| MOTOR_LF | 0.192 |  |  |
| WM_OBK_BODY | 0.187 |  |  |
| WM_OBK_FACE | 0.185 |  |  |

**Supplementary Figure 1.**
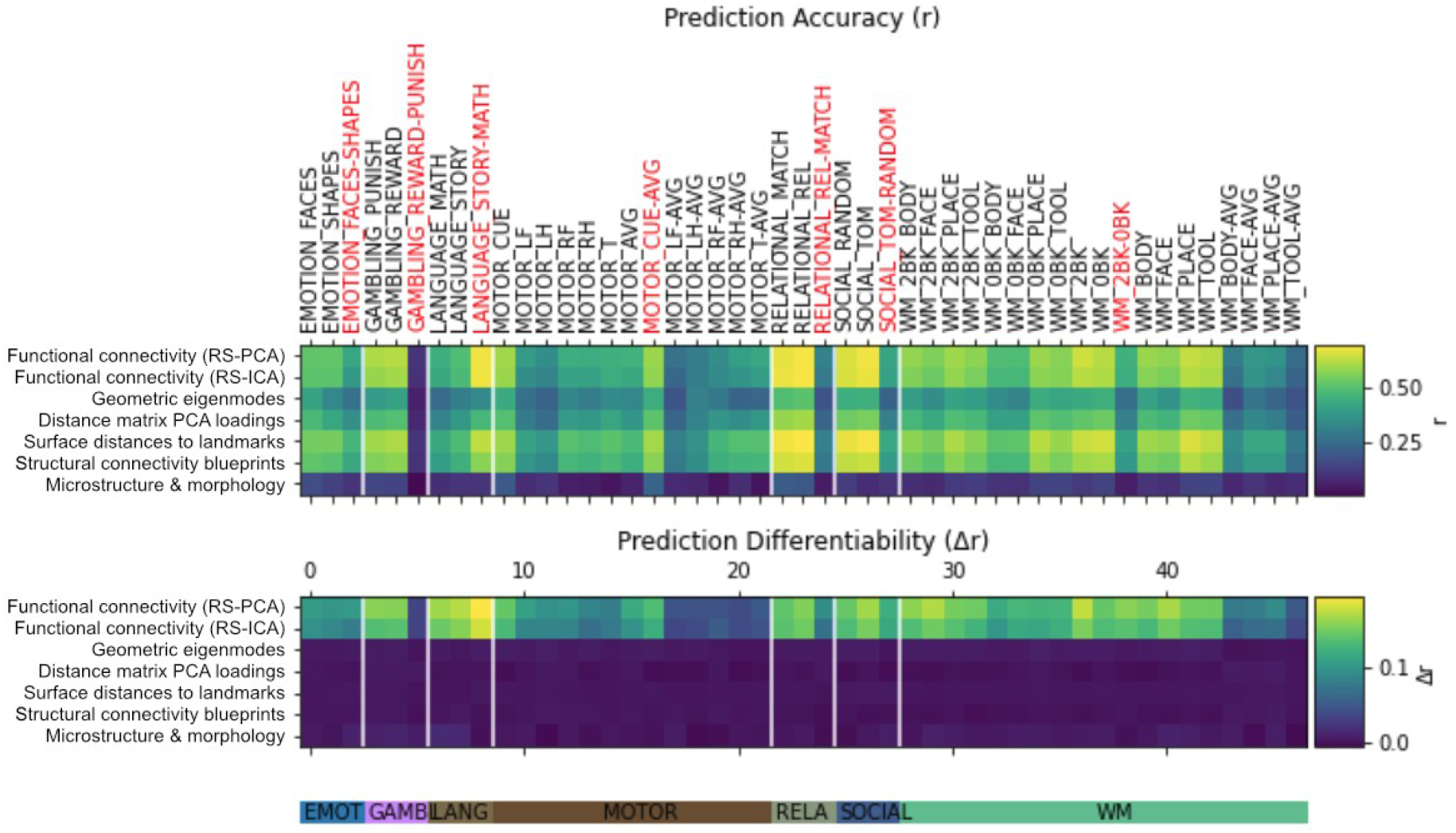
Results for all task contrasts

**Supplementary Figure 2:**
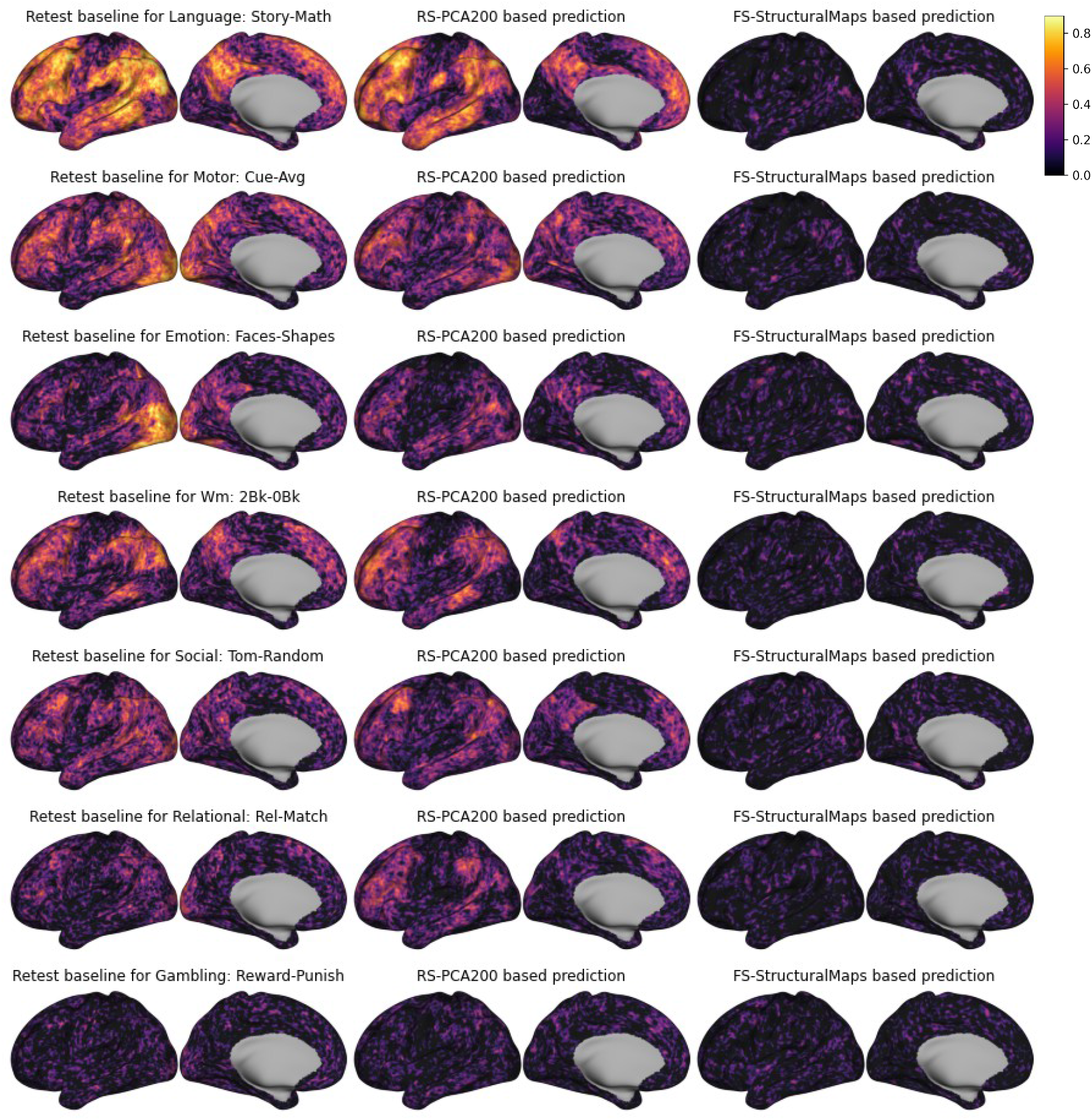
Vertex-wise accuracy for PCA and structural maps

**Supplementary Figure 3:**
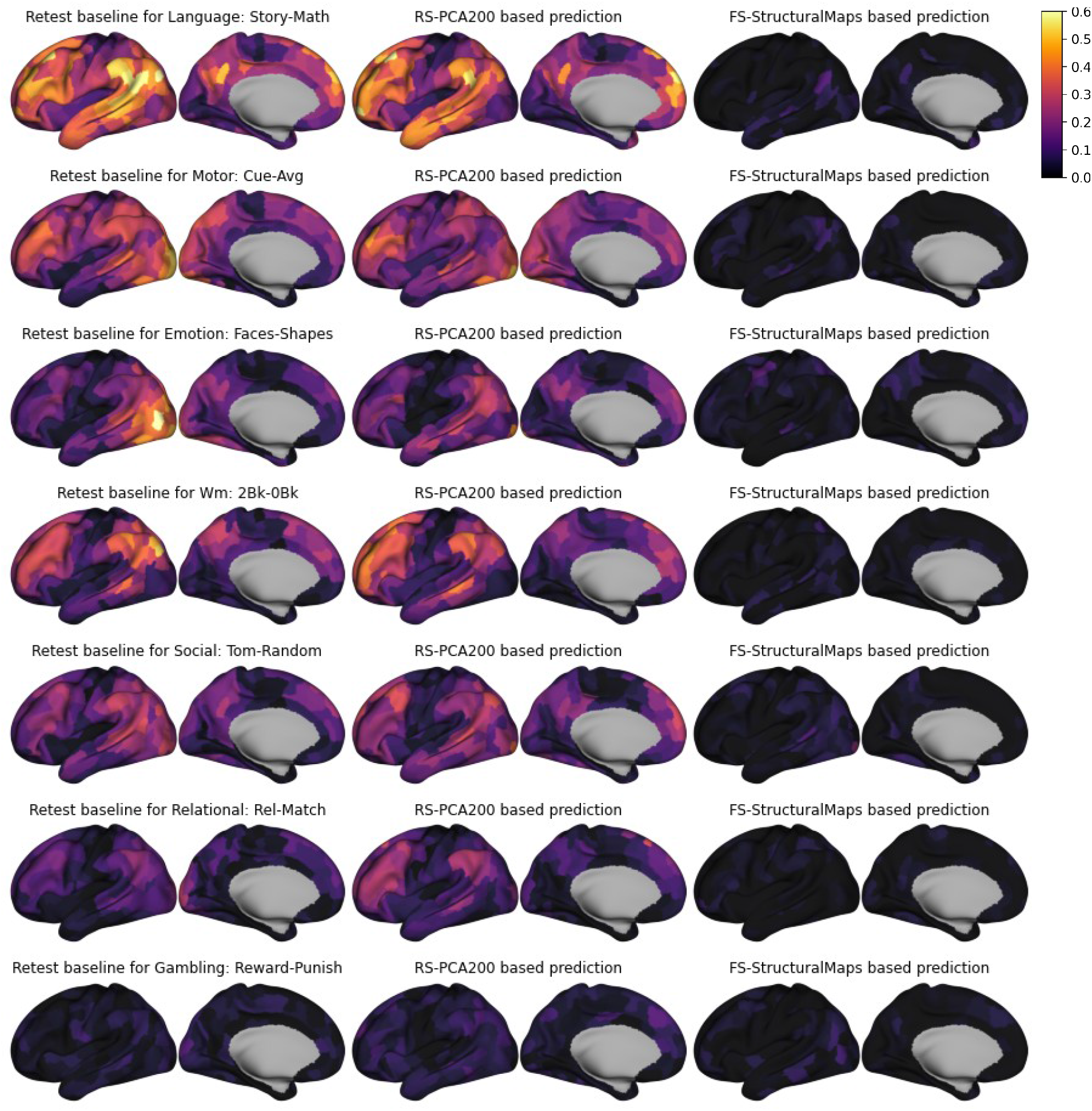
Within-parcel discriminability for PCA and structural maps

**Supplementary Figure 4:**
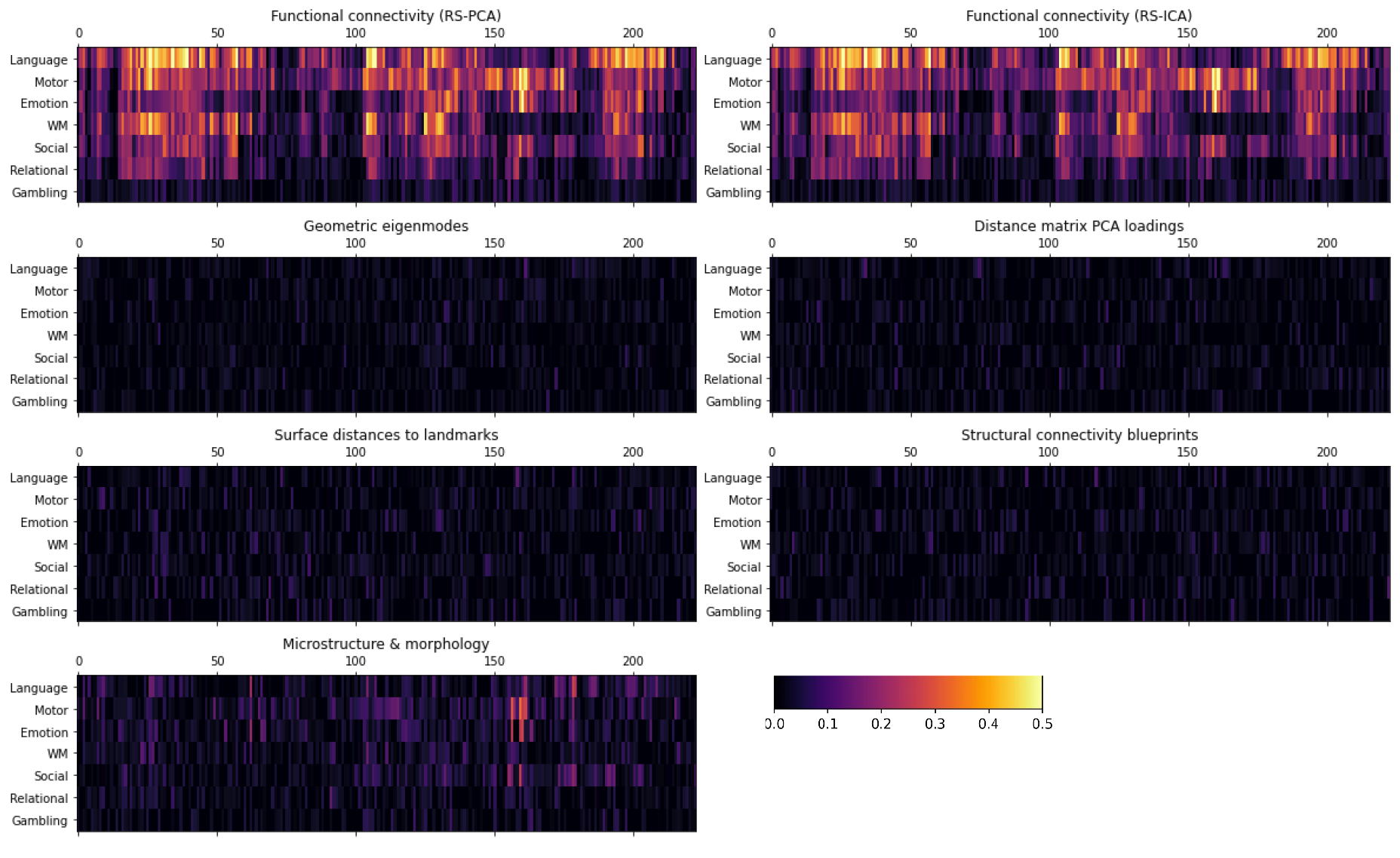
Discriminability scores of the per-parcel models

**Supplementary Figure 5:**
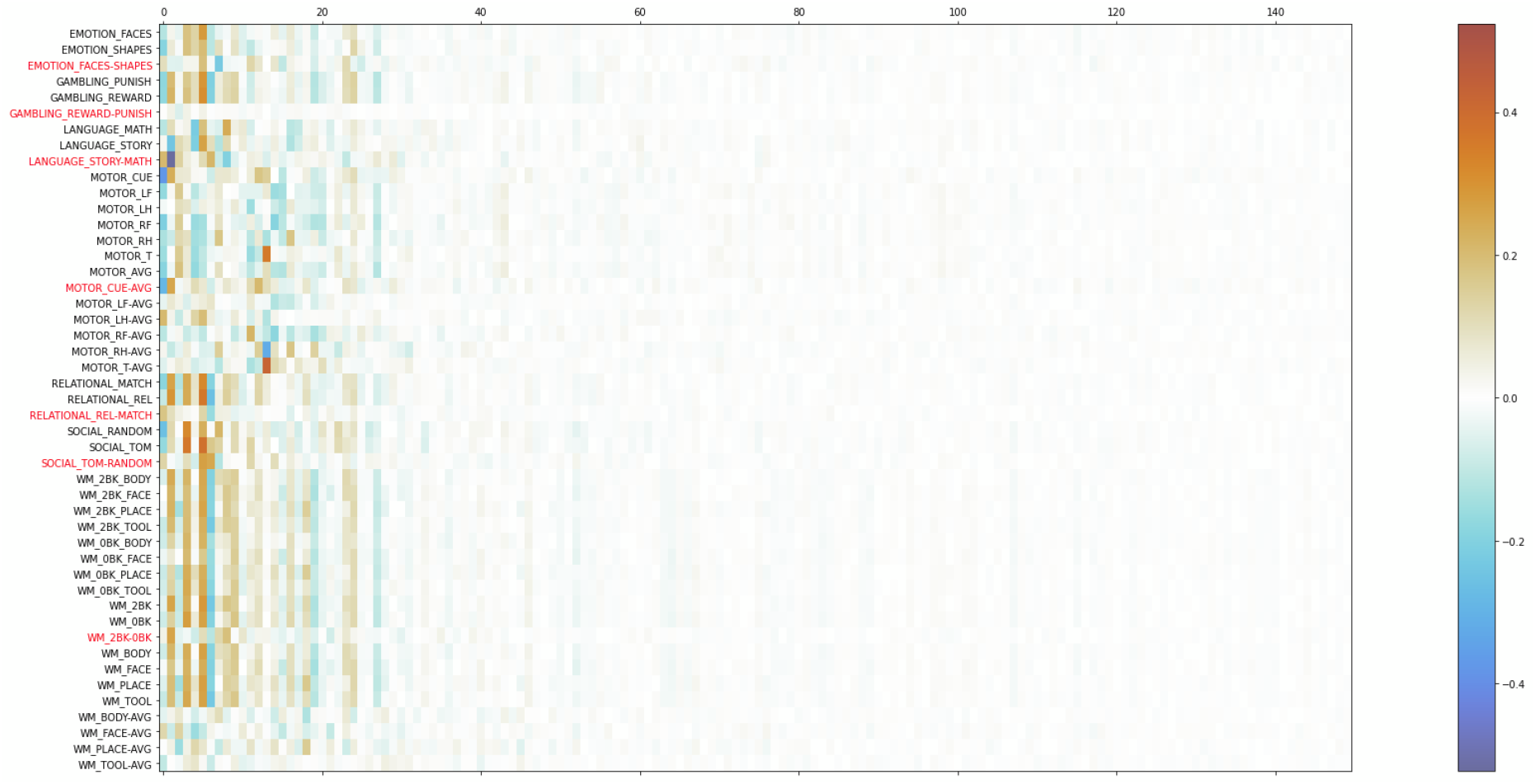
Linear model coefficients or RS-PCA (first 150 of 200 components)

**Supplementary Table 2.**
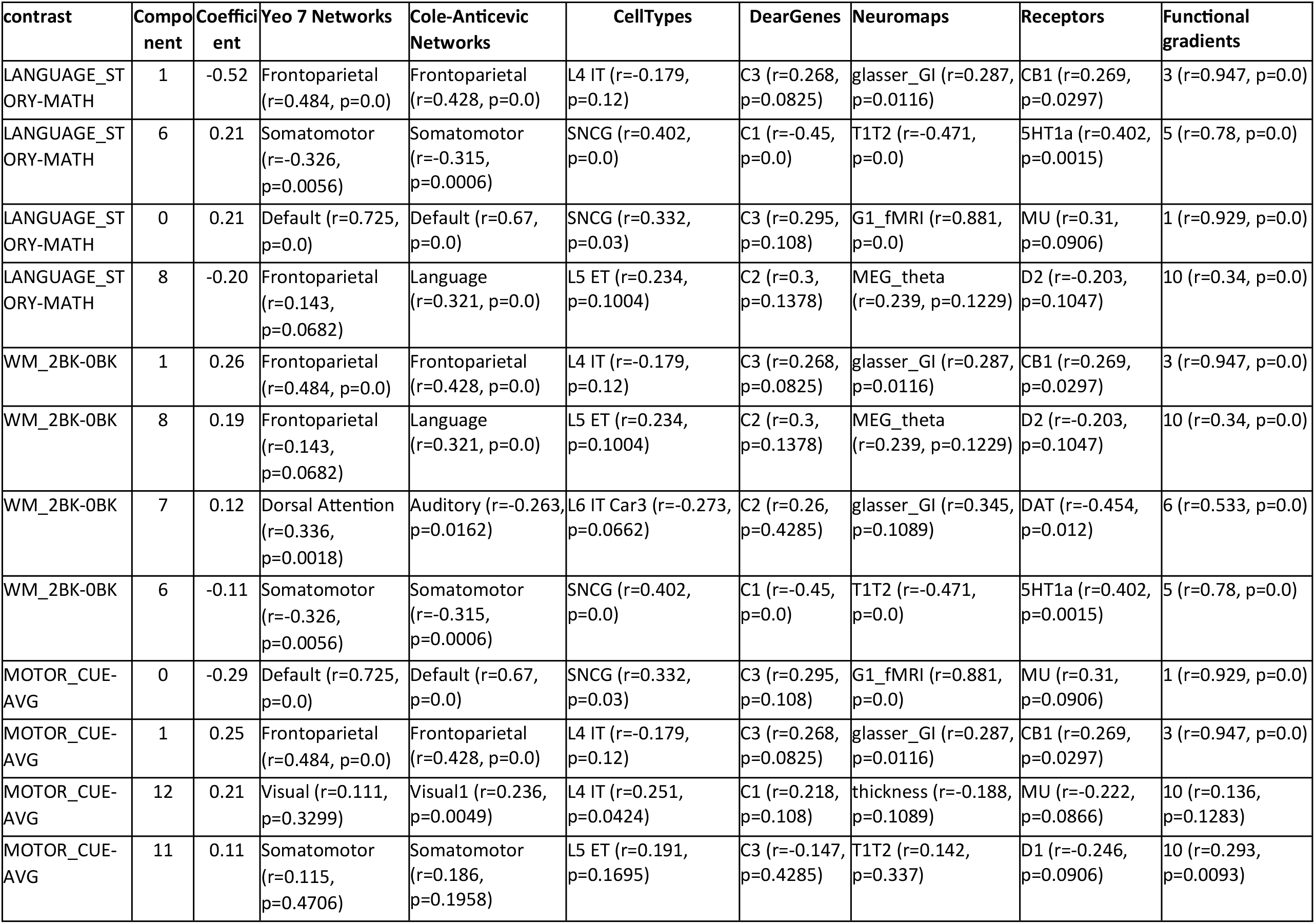
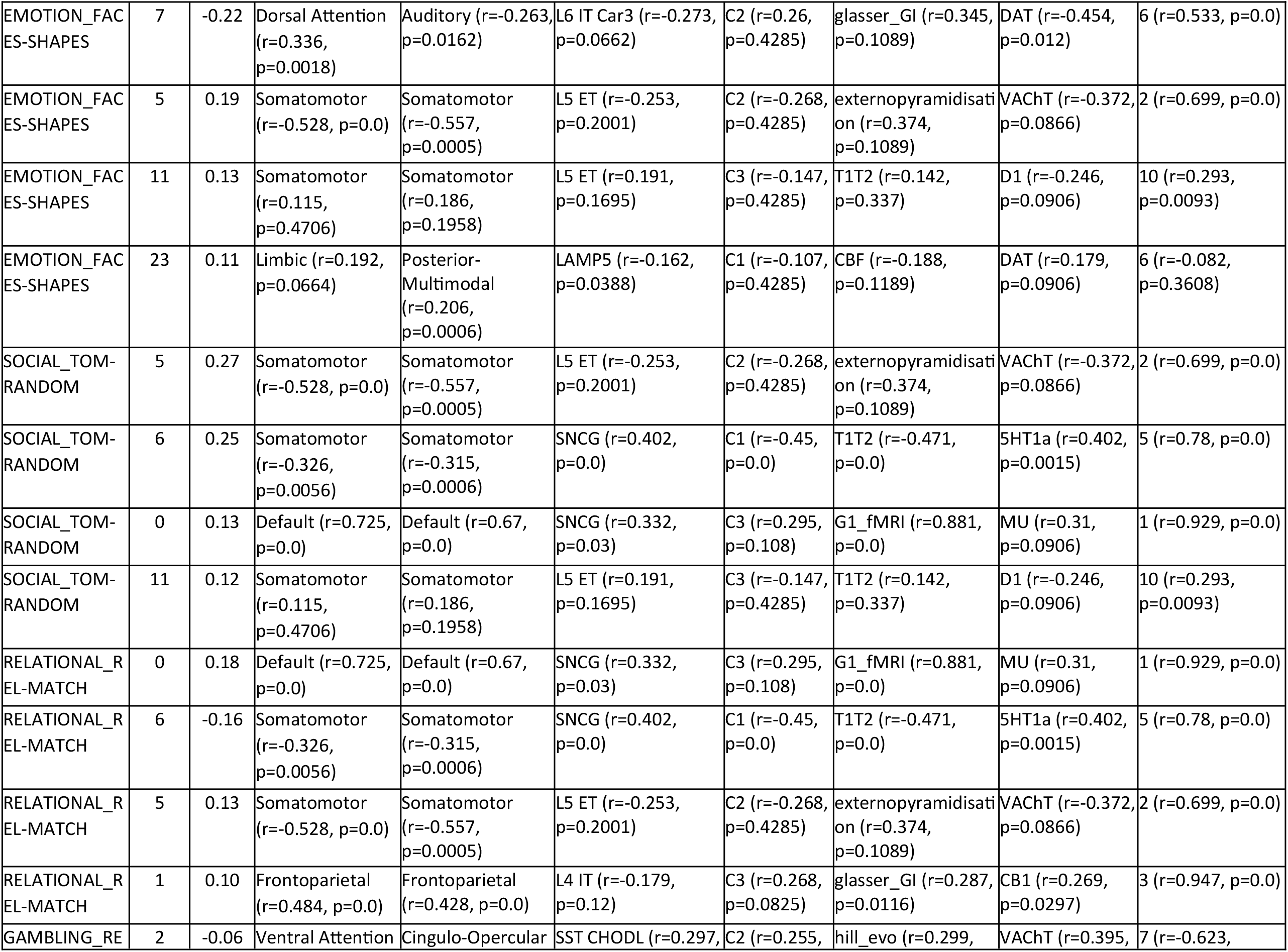
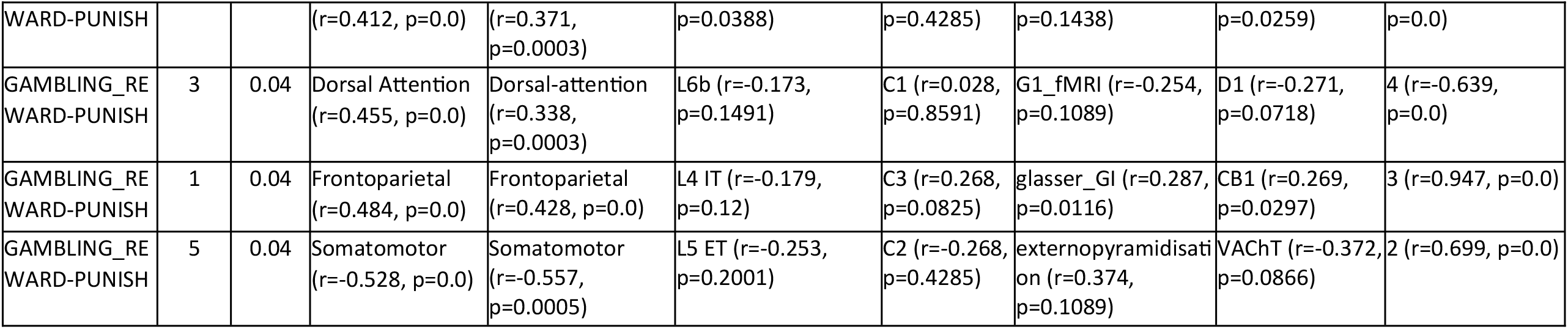
Highest correlating reference map by type for the three most predictive RS components.

**Supplementary Figure 6:**
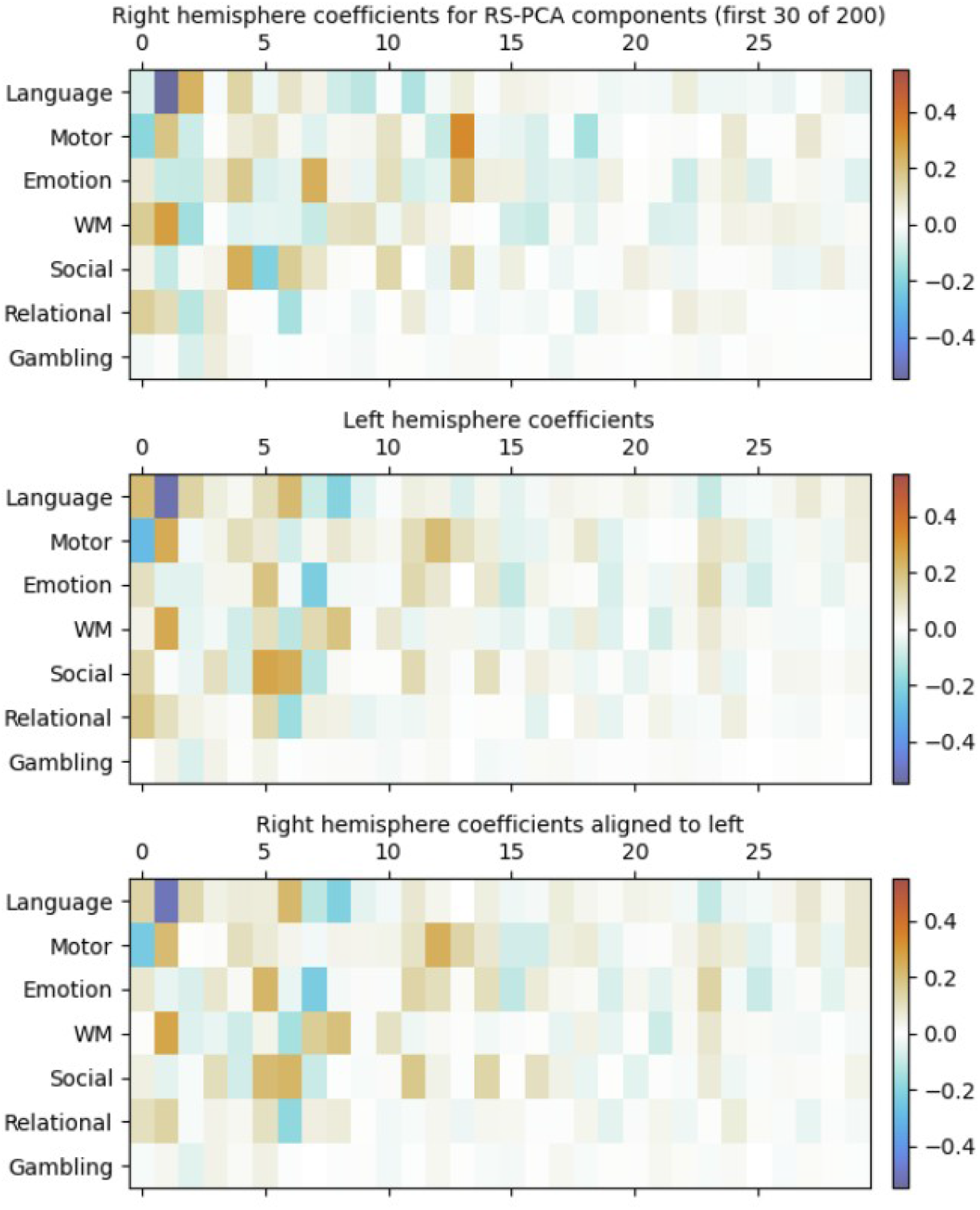
Linear model coefficients compared across left and right-hemispheric RS-PCA models

**Supplementary Figure 7:**
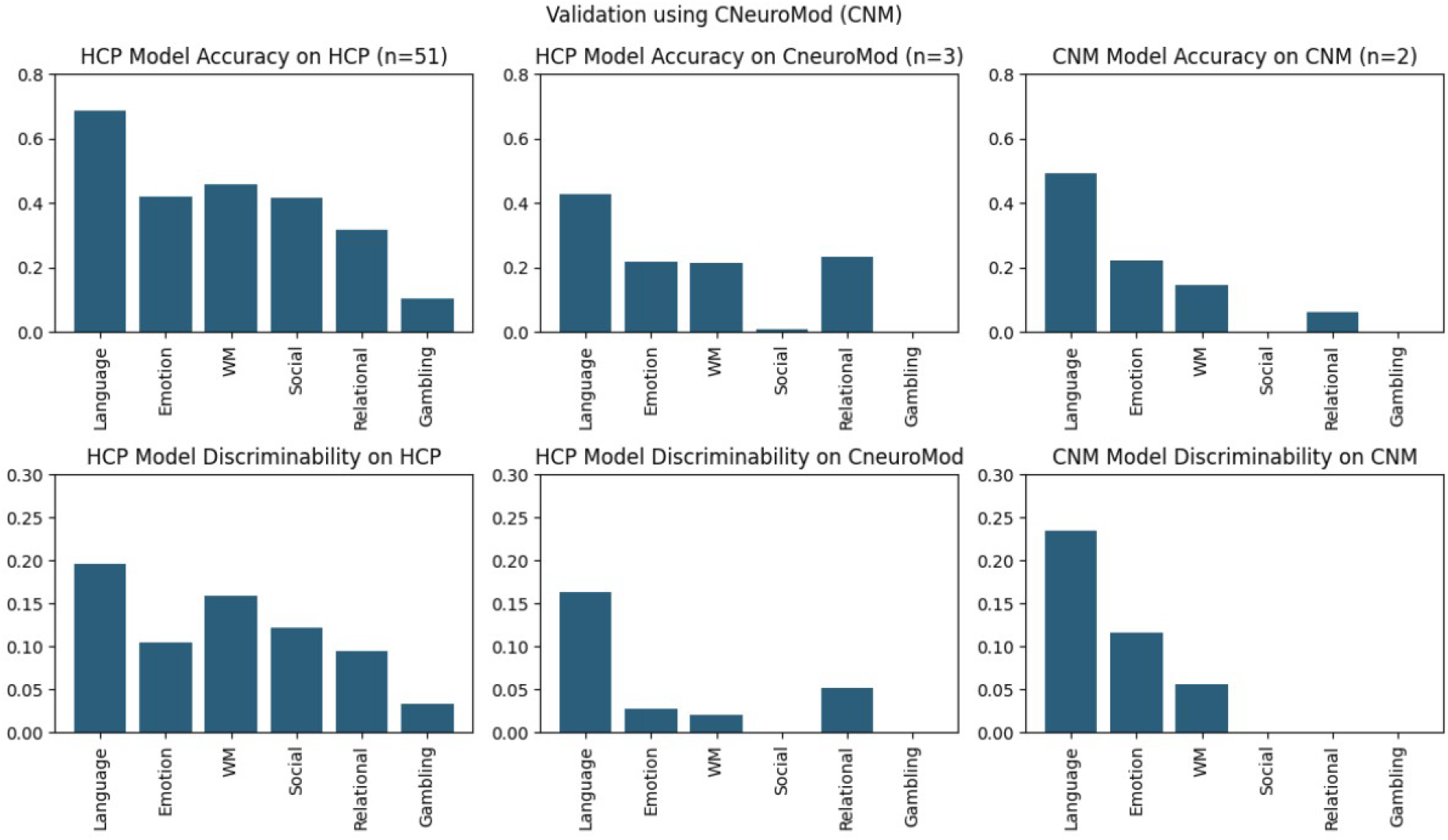
Validation of RS-PCA model using the CNM dataset

**Supplementary Figure 8:**
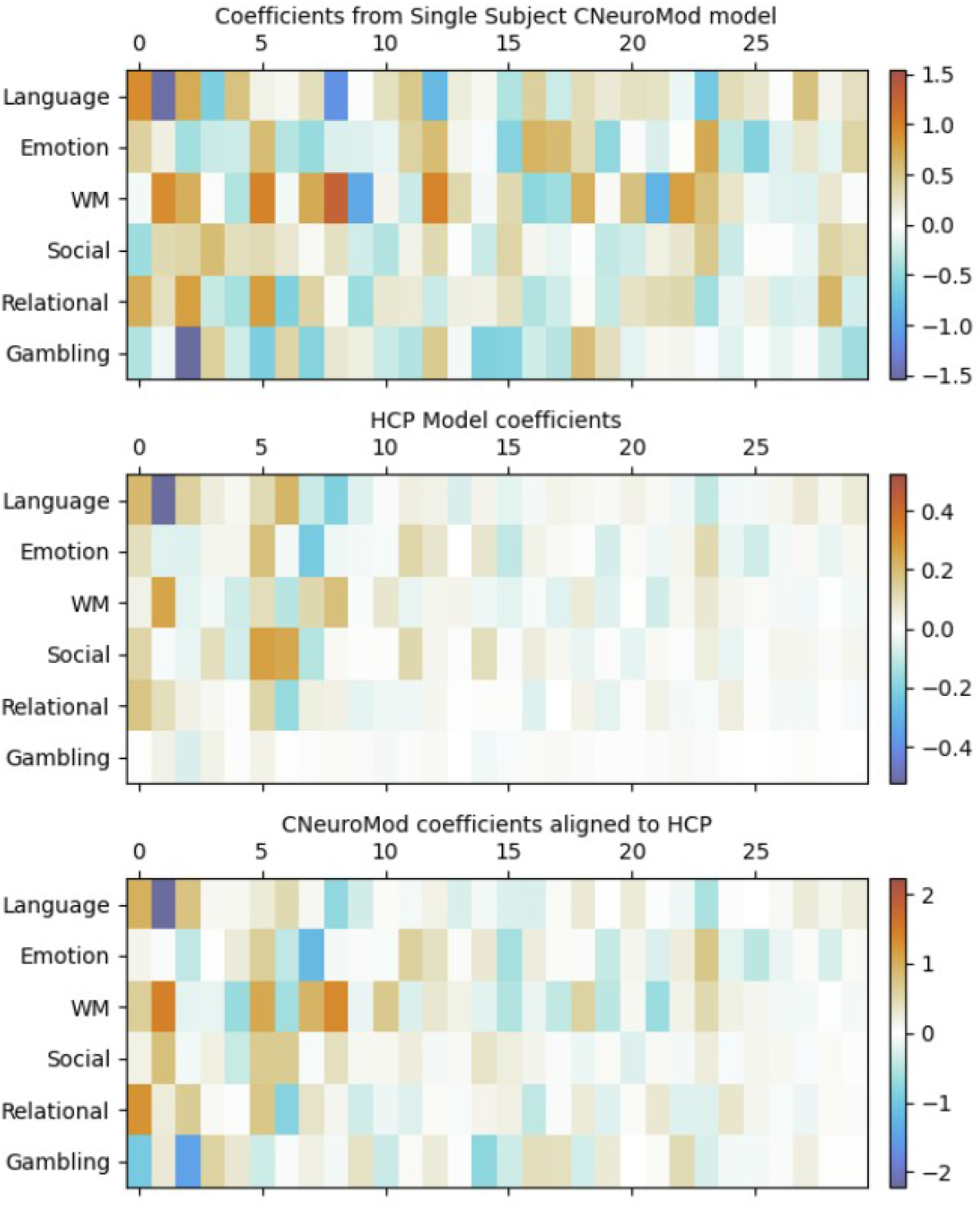
Linear model coefficients compared across RS-PCA models fit on different datasets (HCP and CNM)

**Supplementary Table 3.**
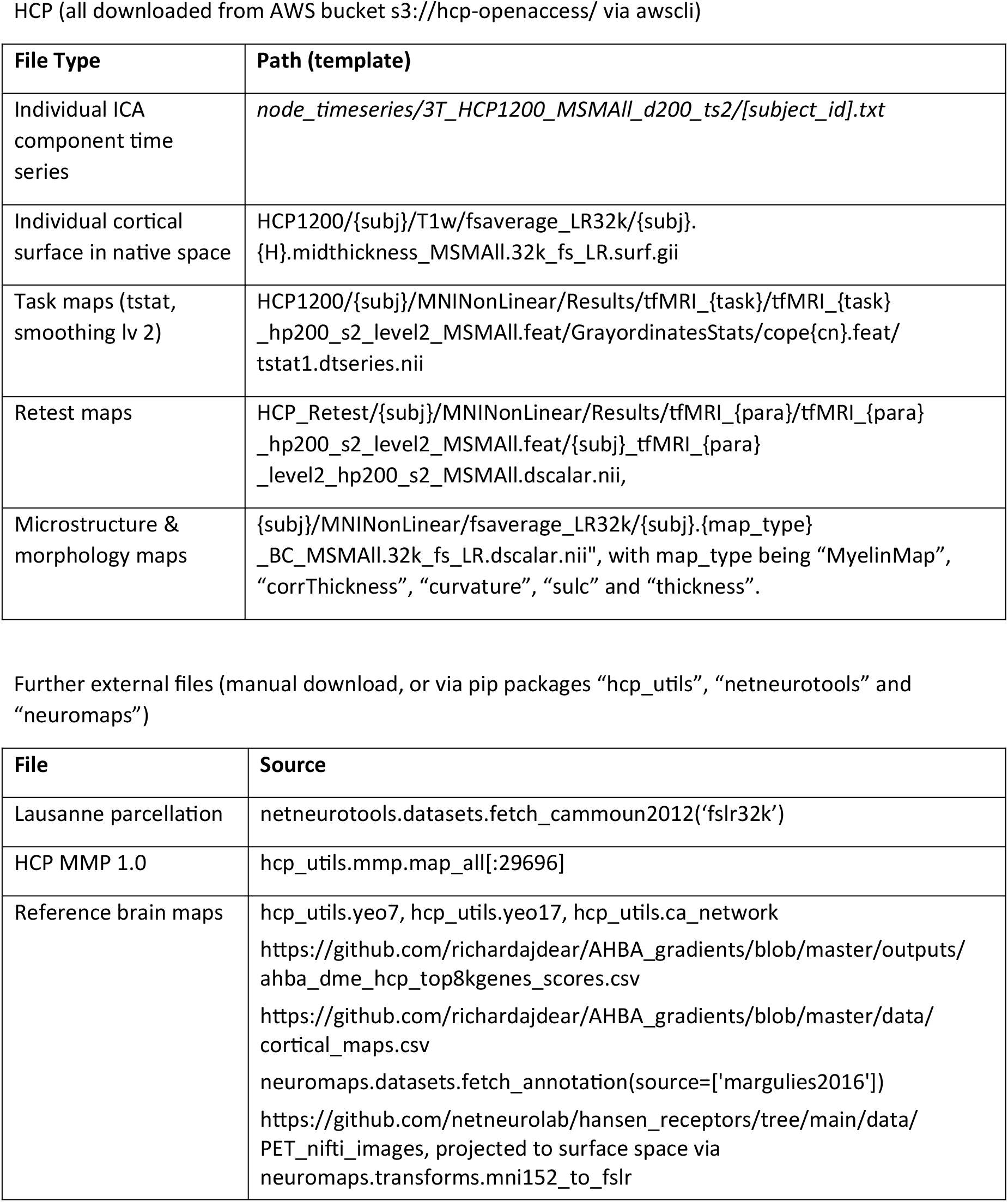
Overview of used external accessible files.

